# Hierarchical and non-hierarchical network flows generate complementary representational dynamics in human visual cortex

**DOI:** 10.64898/2026.04.21.719298

**Authors:** A. Tzalavras, D. E. Osher, C. Cocuzza, L. N. C. Chakravarthula, R. D. Mill, K. L. Peterson, M. W. Cole

**Affiliations:** Center for Molecular & Behavioral Neuroscience, Rutgers University, Newark, NJ, 07102; Graduate Program in Neuroscience PhD Program, Rutgers University, Newark, NJ, 07102; Department of Psychology, The Ohio State University, Columbus, OH, 43210; Department of Psychiatry, Brain Health Institute, Rutgers University, NJ, 08854

**Keywords:** visual system, hierarchy, intrinsic connectivity, activity flow, dimensionality, compression, information flow

## Abstract

Hierarchy is considered a fundamental organizing principle of visual cortex, but its functional implications remain debated given the presence of direct (non-hierarchical) connections. Building on recent advances in measuring direct region-to-region functional connectivity in the human brain, and in using that connectivity (rather than, e.g., visual classification training) to construct deep neural network models, we tested the hypothesis that hierarchical and direct connectivity pathways make distinct contributions to the generation of visual functionality. Detailed measurement of visual functionality, connectivity, and their interaction was achieved using 7T MRI and empirical neural network (ENN) models parameterized by empirical connectivity estimates. The classic V1 to V4 hierarchy was recovered in terms of (i) network distance from V1 along the human brain’s direct region-to-region resting-state functional connectome and (ii) on-task representational transformation distance (visual representation dissimilarity) from V1. *In silico* ENN lesion experiments revealed that hierarchical pathways (V1↔V2↔V3↔V4) reduce the dimensionality of neural representations relative to more rapid and high-dimensional representational contributions from direct pathways (e.g., V1↔V4). These findings reveal distinct but complementary roles of hierarchical and direct pathways in generating cortical functionality.

**Significance Statement:** Hierarchy is a foundational organizing principle of cortex, yet its functional consequences remain unclear because of direct, non-hierarchical connections. The visual system, often portrayed as the clearest example of cortical hierarchy, provides a testbed for dissociating hierarchical and non-hierarchical contributions. Using high-resolution 7T MRI with recent advances in measuring direct region-to-region functional connectivity, we mapped the classic V1 to V4 hierarchy in the human brain. Using empirical neural network (ENN) models parameterized by these empirical connections, we determined that hierarchical pathways (V1↔V2↔V3↔V4) reduce representational dimensionality relative to more rapid, high-dimensional contributions from direct pathways (e.g., V1↔V4). These results reveal complementary hierarchical and direct contributions and establish ENN modeling as a general approach for determining pathway-specific functions throughout the brain.

## Introduction

Vision depends on transforming retinal input into neural representations that support perception and behavior. Classical work on the visual system, together with anatomical studies of interareal connectivity, gave rise to the influential view that visual cortex is organized hierarchically across stages such as V1, V2, V3 and V4 (Felleman & Van Essen, 1991; Hubel & Wiesel, 1968; Rockland & Pandya, 1979). Hierarchy has been defined in multiple ways (Hilgetag & Goulas, 2020), including functional gradients (Margulies et al., 2016), patterns of laminar connections (Felleman & Van Essen, 1991), interareal dynamics (Vezoli et al., 2021), and representational complexity (DiCarlo et al., 2012).

Given this flexibility, arguments have arisen that the visual system may be better described as a *heterarchy,* a combination of hierarchical and non-hierarchical properties (Hawkins et al., 2025; McCulloch, 1945). This issue is especially important in early visual cortex, where direct and reciprocal connections complicate a strictly serial account of processing. Thus, a central question is what distinct functional roles hierarchical and direct pathways play in generating visual function.

To address this question, we used the activity flow framework (Figure 1A), which bases the generation of brain functionality on activity flows – the movement of neural activity over network connections (Cole et al., 2016; Ito et al., 2020a; Yan et al., 2021). This framework provides a natural way to operationalize hierarchy relative to V1 in two complementary ways: *network flow distance,* quantifying how far a region lies from V1 over the empirical connectivity graph when both direct and hierarchical pathways are considered, and *representational distance*, quantifying how much a region’s task-evoked retinotopic representations differ from V1 (Ito et al., 2022).

**Figure 1:**
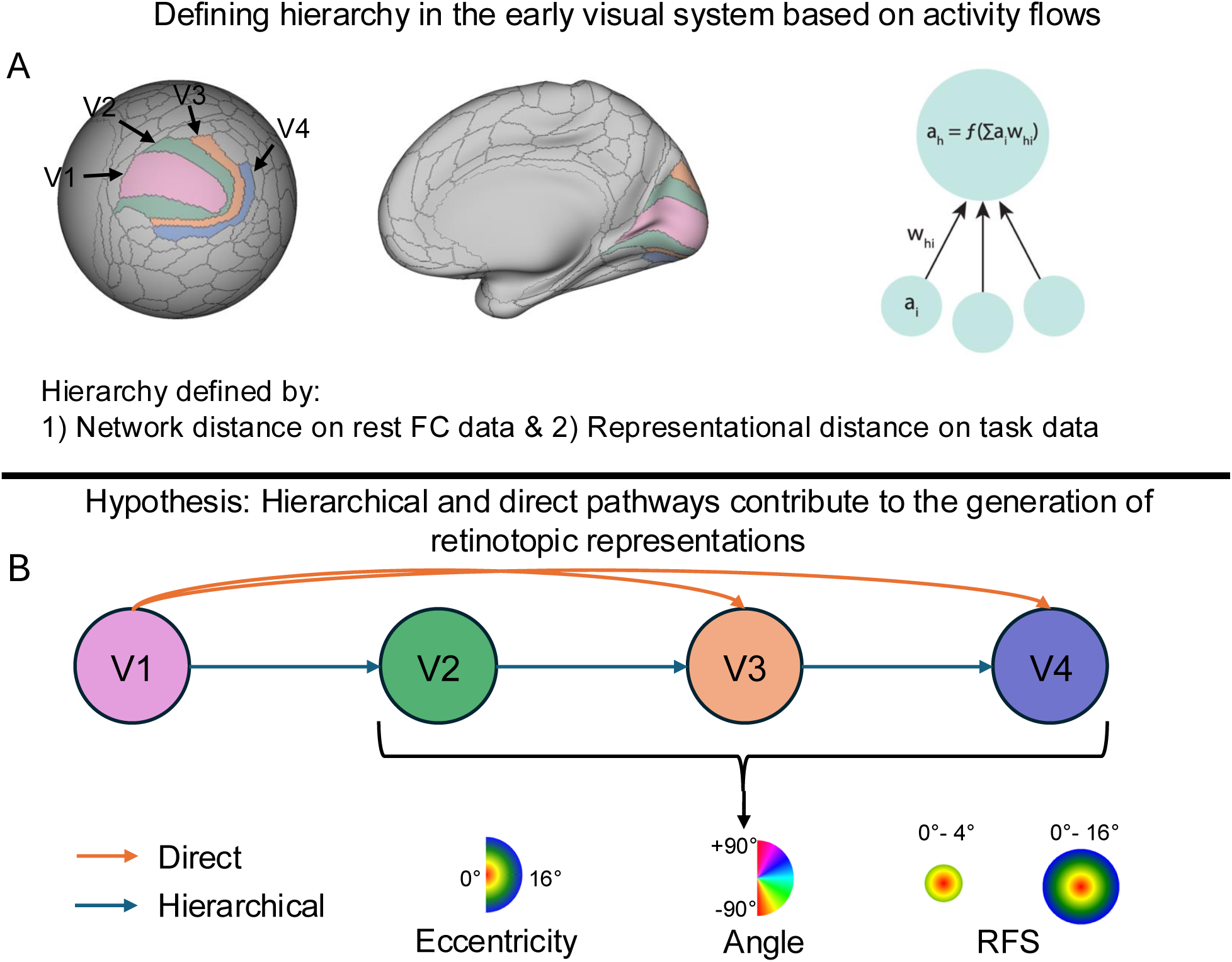
Testing the hypothesis that activity-flow defined hierarchical and direct pathways shape visual functionality in V1-V4. A: Areas V1-V4 depicted on spherical and inflated cortical surfaces. Right, schematic of activity flow framework (Cole et al., 2016): the target activity a_h_ is the weighted sum of source activities a_i_, via connectivity weights w_hi_. In this implementation f is the identity function (no non-linearities). Hierarchy in the early visual system is defined based on resting-state functional connectivity (RSFC) network distance and task data representational distance. B: The inferred hierarchy includes stepwise hierarchical pathways (V1↔V2↔V3↔V4) and direct long-range pathways. Both contribute to visual functionality. Insets illustrate eccentricity, polar angle and receptive field size (RFS) maps. Note that all connections are bidirectional, with directed arrows depicted here to highlight that activity flows originate in V1 (even though they also feedback in subsequent flow steps).

For our network-based measure, we estimated pathway structure from resting-state fMRI data, motivated by prior work showing that resting-state functional connectivity (RSFC) in the visual system is shaped by retinotopic organization (Arcaro et al., 2015; Haak et al., 2018; Heinzle et al., 2011; Watson & Andrews, 2022). We quantified network flow distance using the graph theoretical measurement of communicability (Estrada & Hatano, 2008), which incorporates all connectivity-defined paths to a given node and quantified representational distance from task fMRI retinotopic data. We then tested whether these independent measures converged on the classical hierarchy V1↔V2 V3↔V4.

Building on these hierarchy definitions, we employed ENNs to dissociate the functional contributions of hierarchical and direct pathways. The ENN instantiated activity flow over empirical connectivity using stimulus-evoked V1 activation to generate responses in V2, V3 and V4 (Cole et al., 2016; Ito et al., 2017). Activity propagated over each subject’s empirical connectome and feedback connections were included to reflect the reciprocal coupling known to characterize early visual cortex (Markov et al., 2014). This framework allowed us to test whether downstream retinotopic organization could be generated from V1 activity alone and enabled anatomically interpretable *in silico* lesions. By selectively lesioning direct (V1↔V3, V1↔V4) or hierarchical pathways (V2↔V3, V3↔V4), we quantified their distinct contributions to visual function.

We hypothesized that both pathway types contribute to visual function (Figure 1B), but in distinct ways: hierarchical pathways decrease representational dimensionality, through greater integration across processing stages, whereas direct pathways retain more high-dimensional visual information from V1. Lastly, we hypothesized that pathway dependence would vary with receptive-field scale building on prior non-human primate work showing that direct V1 to V4 connections are biased towards foveal representations, (Nakamura et al., 1993; Yukie & Iwai, 1985). More specifically, we hypothesized that representations with a receptive field large enough to include the visual periphery would rely more on the hierarchical pathways. In contrast, representations with receptive fields small enough to be dominated by the fovea would rely more on the direct pathways.

In summary, we validated two independent measures of hierarchy in the early visual system and used them to delineate the functional contributions of hierarchical and direct pathways in terms of representational dimensionality and across receptive field scales (Figure 1).

## Methods

### Overview

We utilized the Human Connectome Project (HCP) Young Adult minimally preprocessed, ICA-FIX-denoised 7T fMRI dataset, including retinotopy and resting-state acquisitions. After estimating FC, we applied a V1-initiated ENN (a multi-step activity flow model) to the cortical vertex time series. The resulting predictions were then input into a population receptive field (pRF) model (Benson et al., 2018), and the outcomes were compared to the original. We defined hierarchy using two independent metrics: 1) network flow distance from V1 based on RSFC and 2) representational distance from V1 based on task/retinotopic representations. We assessed hierarchical and direct contributions to visual functionality by performing *in silico* lesions on our V1-initialized ENN. We calculated dimensionality using the participation ratio (detailed below) and we tested for links between the hierarchical pathways and large receptive field size representations (Fig. 2).

**Figure 2:**
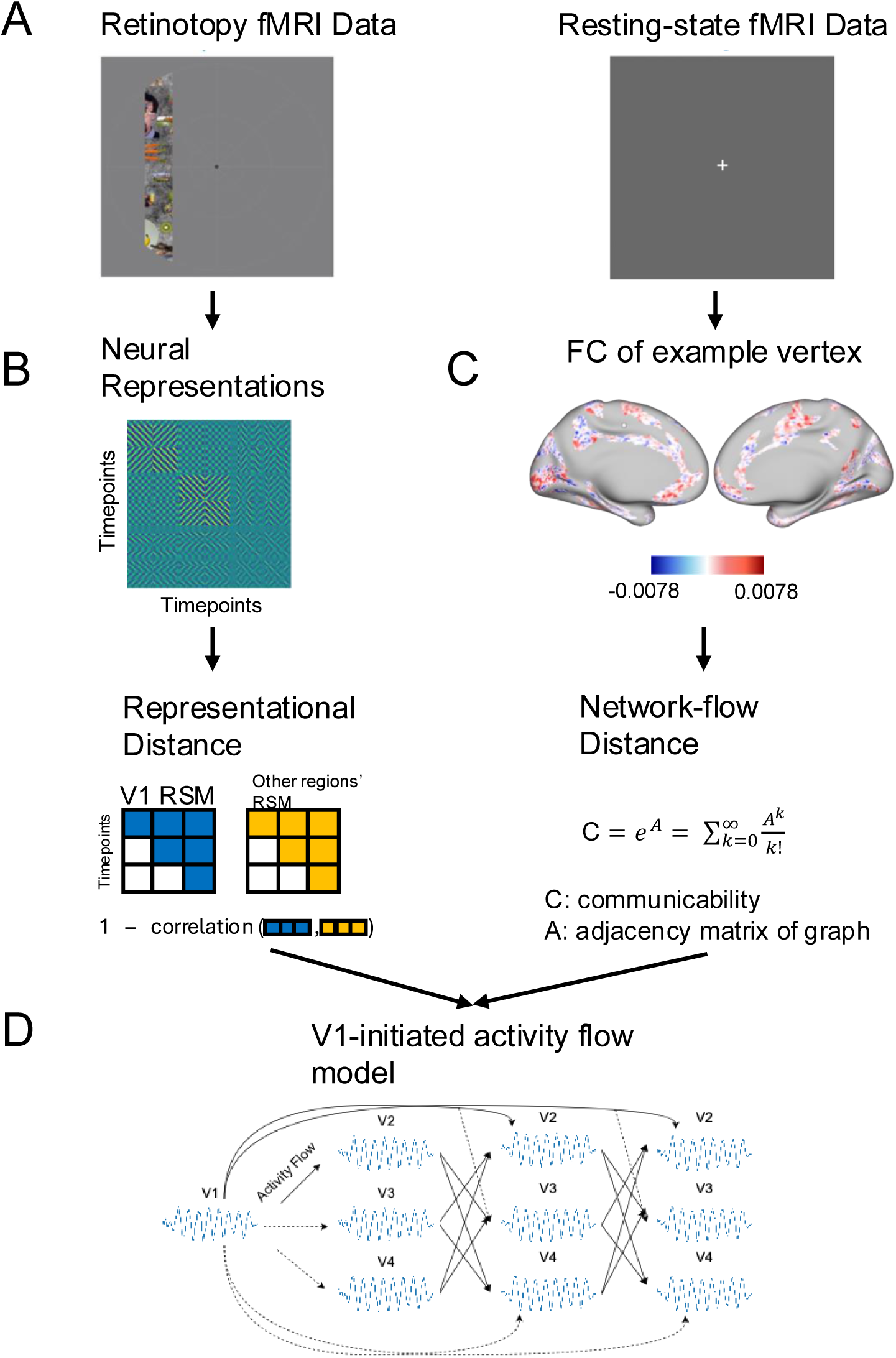
Analysis pipeline and hypotheses linking hierarchy to retinotopy. A: 7T Human Connectome Project fMRI rest and retinotopy data from 40 participants (6 runs with rotating wedges, expanding/contracting rings and moving bars, 4 resting-state runs ∼ 1 hour, Benson et al., 2018). B: Top: Region-wise neural representations summarized as representational similarity matrices (RSMs). Bottom: Representational distance from V1 to each region, defined as 1 – correlation between their RSMs, used as the first hierarchy metric. C: Top: Whole cortex functional connectivity (FC) of an example vertex (see Method and Fig. 3). Bottom: Network flow distance, computed via communicability, used as a second hierarchy metric (see Methods and Fig. 4). D: V1-initiated activity-flow model (after Cocuzza et al., 2024) propagating activity from V1 to downstream regions (V2-V4) via FC weights.

### Data collection and preprocessing

We used data from 181 healthy young adults from the HCP Young Adult 7T retinotopy dataset (Benson et al., 2018), part of the WU-Minn HCP Consortium (Van Essen et al., 2013). Out of the 181, 40 subjects were selected at random (20 females). Participants underwent imaging on a customized 7T Siemens Magnetom scanner, with T1-weighted and T2-weighted structural scans at 0.7-mm isotropic resolution and six retinotopic fMRI runs totaling 30 minutes (1,800 time points) at 1.6-mm isotropic resolution and 1-second TR (TE = 22.2 ms). Four runs of resting-state data (∼1h of data) with the same parameters were also collected. Data were minimally preprocessed using the HCP pipelines (Glasser et al., 2013) and denoised using ICA-FIX (Glasser et al., 2018). All participants provided informed consent, and data were deidentified prior to public release. We used the spatially down-sampled version of the data resulting in 2-mm spatial resolution. Finally, for structural connectivity (SC), we used HCP Young Adult 3T diffusion MRI acquisitions collected with a three-shell protocol (b=1000,2000,3000 s/mm^2^) at 1.25-mm isotropic resolution (TR = 5520ms, TE = 89.5 ms) across 6 runs (both phase-encoding polarities), as described in the HCP protocol.

### Retinotopic mapping

One prominent way to elucidate the retinotopic organization of the human visual cortex is pRF modeling (Dumoulin & Wandell, 2008). The pRF model treats each voxel/vertex BOLD time series as reflecting the pooled responses of neurons within that voxel/vertex, whose combined spatial sensitivity is approximated by a single population receptive field. Also, in a common formulation, this spatial sensitivity can be approximated by an isotropic (circularly symmetric) 2D Gaussian. In the compressive spatial summation model used here, the Gaussian’s center and size are estimated by predicting the BOLD time series from the stimulus sequence. Stimulus contrast within the Gaussian is pooled and then passed through a compressive power law with parameters chosen to best fit the measured data. (Benson et al., 2018; Kay et al., 2013). The stimuli presented during the experiment include contiguously moving apertures constrained to a circular region with diameter of 16 degrees, as the subject fixates on a central point. The outputs of the model include the preferred polar angle (angular position of the Gaussian with respect to the point of fixation), eccentricity (distance from fixation in degrees) and receptive field size (Gaussian spread in degrees) of each voxel/vertex as represented by the optimal parameters of the fitted Gaussian. In this study we used the retinotopic dataset and process employed by (Benson et al., 2018) to fit the pRF model. This process involves a nonlinear optimizer solving the model for each vertex separately. We provided the ENN-generated and original time series to the pRF model and then statistically compared the results to estimate model accuracy.

### Activity flow mapping

Activity flow mapping has been used in over a dozen studies (e.g., Cole et al., 2021; Ito et al., 2020; Mill et al., 2017), each demonstrating its ability to provide a mechanistic explanation of the generation of neural activity under empirical constraints. Activity flow mapping simulates the propagation of neural activity – and the information it carries – throughout brain networks. By forcing our model to initiate flows in V1, we add additional causal validity by simulating the path that visual information travels as it enters cortex. The resulting ENN model was V1-initiated and multi-step, in the sense that activity propagated from V1 over multiple simulated time steps, with each subject’s individualized resting-state network organization constraining those flows.

Each subject’s individualized ENN model was constructed with a focus on the early visual system (V1-V4), using empirical connectivity weights at the vertex level. We used V1-V4 connections retained after parcel-level masking by the group-average SC matrix, which constrained the set of allowable flows, while ENN weights remained vertex level FC estimates. We did not use SC edge weights in our ENN models because, while SC reflects axon bundle projections, in contrast to FC they do not reflect the many additional biological features that influence the transmission of neural signals (e.g., the aggregate effects of synaptic weights). The SC matrix was computed following the same methods described in Peterson and colleagues (2025). Briefly, we estimated parcel-level SC from HCP Young Adult 3T diffusion MRI using the HCP diffusion preprocessing pipeline, followed by DSI Studio deterministic tractography with generalized Q-sampling imaging. Edge weights were defined as normalized streamline counts between parcels, and the group-average SC matrix was based on 352 subjects. We focused on early visual areas because the retinotopic pRF model was more successful in explaining the observed neural activation in those areas, allowing us to focus on retinotopy. The initial model-generated activation of each vertex in each target area (V2-V4) was produced using the activity flows from source vertices in V1 to each of the target vertices. The activity flows were computed according to the activity flow framework as source neuronal activations multiplied by the intrinsic connectivity weights between source and target (Fig. 2). The following simulated flow steps generated the activation of each target vertex by using as sources the generated activation of the targets from the previous step, along with flows from sustained activity in V1 (modeling the continuous visual input during retinotopic mapping). Lastly, to incorporate an abstract temporal component in the generation of activations, we repeated this process for 6 steps (see Figure 2). The number of flow steps was determined as the step wherein there is an increase of less than 1% in the explained variance (R^2^) between the generated and actual time series, averaged across subjects. This R^2^ measure was based on time series calculated as the average across vertices and across regions for both the generated and actual signals among V2, V3, and V4 (both hemispheres). Our model can be characterized in terms of equations as follows: Let 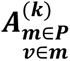 be the activation of the v-vertex that belongs to the m-parcel at the k-th step of our model, with P being the set of target parcels of our model. P = {V2, V3, V4}. Then our model can be defined iteratively as:

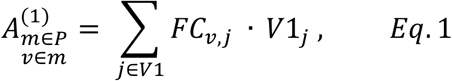

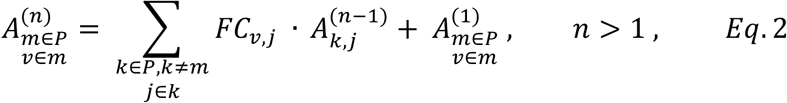

Note that V1 activity was maintained across multiple steps due to the extended nature of visual stimulation in the HCP 7T retinotopic dataset used here. Rather than visual stimulation being transient relative to the hypothesized flow steps (expected to be between 10 ms and 100 ms per step), it was expected that visual stimulation (i.e., V1-initiated flows) would have been extended across all flow steps simulated here.

### FC estimation

We developed a novel method to calculate RSFC by combining graphical lasso (Glasso) regularization with PC regression (Fig. 3). Our motivation was to develop a method that 1) benefits from the substantial reduction of causal confounds in partial correlation and multiple regression FC measures relative to pairwise Pearson correlation (the field standard) (Peterson et al., 2025), 2) increasing the repeat reliability of FC estimates using regularization (Peterson et al., 2025), and 3) applying these computationally-intensive approaches developed with parcel-level FC to vertex level data for the first time. This graph constrained principal component regression (GC-PCR) approach involved two steps of FC calculation: First at the parcel level and second at the vertex level. Initially we calculated parcel-level FC using Glasso, a regularized partial correlation approach previously shown to reduce FC confounds (e.g., a third parcel inducing a false connection between two downstream parcels) while reducing low reliability from overfitting to noise (Peterson et al., 2025). Parcels were defined based on the Glasser parcellation (Glasser et al., 2016). Each vertex of the resting-state data was z-scored prior to averaging across vertices. Publicly released toolbox code was used to estimate Glasso at the parcel level (https://colelab.github.io/ActflowToolbox/). The lambda hyperparameter of the Glasso algorithm, determining the degree of regularization was estimated by grid search over a range of 20 values (10^-7^-10^2^, log-spaced), for each subject, with 10-fold cross validation. This selected the lambda parameter yielding the highest accuracy (R^2^) on predicting held-out test data. Vertex-to-vertex connections that belonged to parcels that were disconnected in the parcel-level connectivity matrix were assigned a weight of 0. Thus, Glasso served as a type of filter, removing from each target vertex all the vertex level connections that belonged to disconnected parcels.

**Figure 3:**
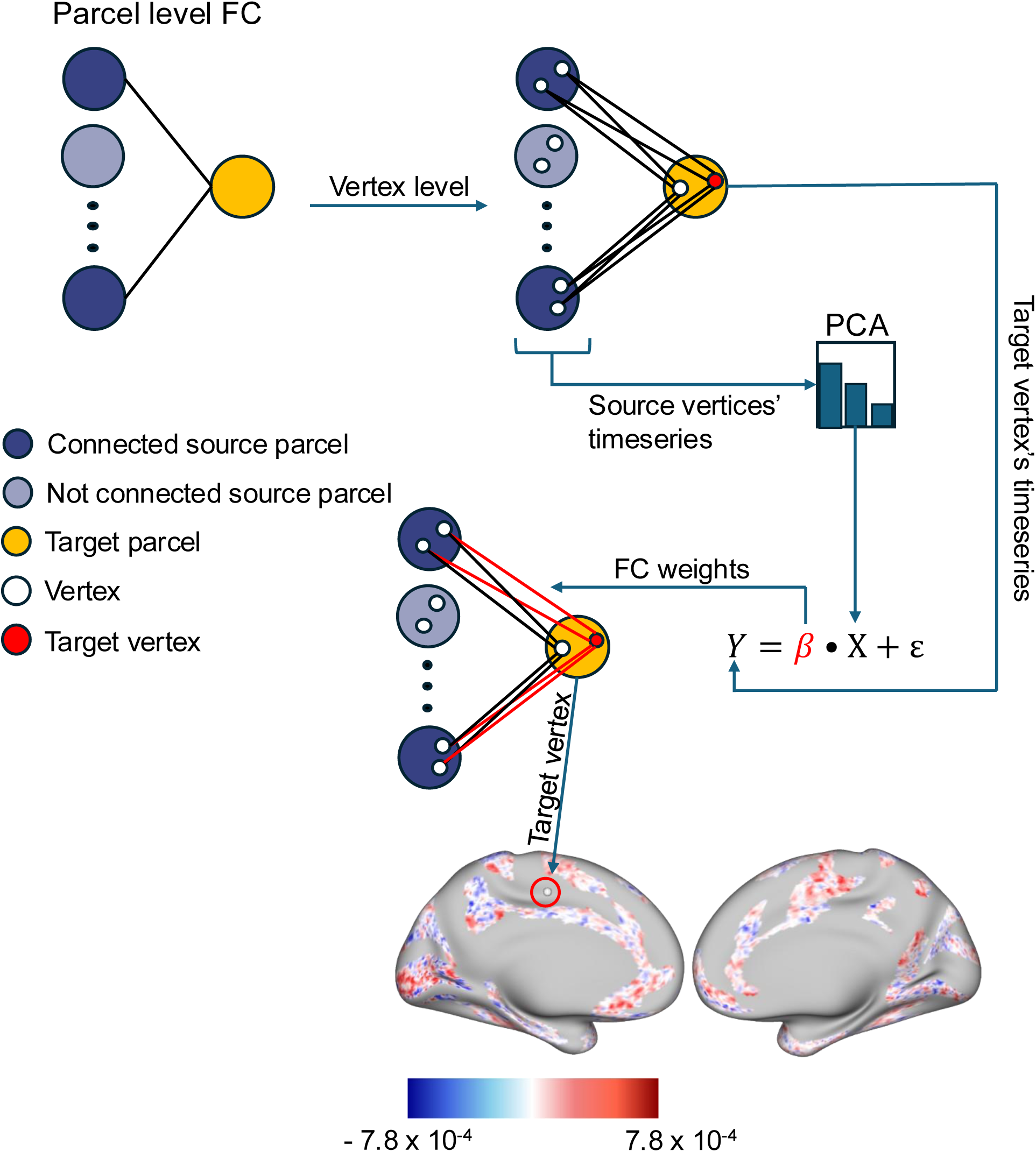
Dense, vertex-wise functional connectome via graph constrained principal component regression (GC-PCR). Parcel-level FC derived from graphical lasso (Glasso) (see Methods) identifies source parcels connected to a target parcel. This reduces confounding of functional connections to estimate direct connections between parcels, with lasso regularization reducing overfitting to noise and effects of multicollinearity. Moving to the vertex level, the time series of the source vertices within connected source parcels are dimensionality-reduced with PCA, which (like lasso) is a form of regularization that reduces overfitting to noise and effects of multicollinearity but, unlike lasso, also reduces computational demands and is unlikely to set any connections to 0. For each target vertex, its time series are predicted from a linear regression model where predictors are the dimensionality-reduced time series of the source vertices. The resulting regression weights β, transformed back to the original vertex space, provide vertex-wise FC from the source vertices to the target vertex. An example cortical map is shown for a target vertex (warm/cool colors indicate positive/negative weights respectively). Iterating this process for each target vertex of each target parcel yields the dense connectome. While the obtained connection estimates were undirected, we interpreted them as bidirectional and equally weighted in both directions, given evidence that most cortico-cortical connections are bidirectional and equally weighted in non-human primates (Markov et al., 2014). It will be important for future work to thoroughly validate methods for estimating directed/effective connectivity in the human brain at the voxel/vertex level.

For each target vertex, the vertices that survived parcel-level filtering were used as sources in a multiple regression model. As is standard in PC regression, the regression took place on the PCs calculated from the time series across all source vertices for that target vertex. The number of retained PCs was determined using a two-step cross-validation procedure, testing for the ability to predict held-out resting-state fMRI data in target vertices. This procedure started from the first component all the way to the maximum number of PCs (minimum between the number of datapoints - 1 or number of features - 1) with a step of 100 PCs. After the optimal number was identified, the second step initiated. Here, a finer search with a step size of 10 was applied with the number of components ranging from the optimal number from the previous step – 50 to the optimal number from the previous step + 50. For each PC number, 10-fold cross validation was performed on the target’s time series and the mean squared error (MSE) was calculated for each fold. The average MSE across folds was calculated and this number represented the accuracy of the prediction for that number of components. The number of components with minimum MSE was selected. After this selection, multiple linear regression was performed using the optimal number of PCs. The resulting weights were transformed back to the original ∼ 64k vertex space, as is standard in PC regression. Functional connectivity of example vertices was visualized using Connectome Workbench (Marcus et al., 2011).

### Retinotopic receptive field estimation

After generating the time series for each vertex of the early visual system, we extracted the time series from the last step and provided them to the pRF model to generate retinotopic receptive field estimates (eccentricity, angle and receptive field size). These estimates were then compared against the actual ones for each vertex using linear correlation for eccentricity and receptive field size and circular correlation for angle. Spearman correlations between the actual and generated eccentricity and receptive field size across vertices for each region for each subject were computed. Statistical significance was estimated using a one-sample t-test against 0 for each region across subjects, Bonferroni-corrected for multiple comparisons. The same approach was followed when time series provided to the pRF model were the results from the hierarchical and the direct *in silico* lesions experiments.

### Statistics on the generated time series

For assessing the prediction accuracy of the model-generated time series, we used the MaxT nonparametric permutation testing approach to correct for multiple comparisons (Nichols & Holmes, 2002). Permutation testing is thought to reduce the often overly-conservative multiple comparison correction of Bonferroni, while also reducing the often overly-liberal multiple comparison correction of false discovery rate (Eklund et al., 2016). This involved 1000 permutations of the FC matrix of each subject, re-running the model, computing per vertex R^2^ at the final step and averaging across runs and subjects. For each permutation we recorded the maximum R^2^ across the 2,565 vertices of V2-V4 for both hemispheres, yielding a null distribution of per-permutation maxima. For each vertex, the model’s R^2^ (identically calculated) was compared to this distribution in order to compute nonparametric p-values that were family-wise error corrected for multiple comparisons. This was used to assess statistical significance for the generated time series at the vertex level.

We also assessed the prediction accuracy of the generated time series at the parcel level. We followed a similar analysis as described above based on the MaxT nonparametric permutation test. This time we averaged the R^2^ values obtained from the null models across vertices for a given parcel (also averaging across runs and subjects as described above). This resulted in a single R^2^ value for each parcel for each of our null models. For each permutation, we recorded the maximum R^2^ among the 6 parcels, creating our null distribution. The average R^2^ value from our model (identically calculated) was compared to this distribution.

Lastly as a sanity check, we report the average across runs R^2^ values based on the averaged time series across vertices for each parcel. To assess statistical significance, we used a one-sample, one-sided Wilcoxon ranked test across the R^2^ values across subjects for each parcel and then Bonferroni corrected for multiple comparisons. This approach is more forgiving, despite the conservative nature of Bonferroni correction, since each parcel’s signal-to-noise ratio was increased by averaging. Statistics for both the parcel and the vertex level were run on the 6^th^ step of the model.

### Network flow distance

The first criterion we used to establish a hierarchy in the early visual system was based on network flow distance (Fig. 4). By network flow distance, we mean the ease of communication (flow) between regions in our FC graph. Notably, our use of partial correlation FC, rather than the field-standard Pearson correlation FC, made network flow distance more meaningful, as partial correlation FC reduced the number of confounding “shortcuts” that would invalidate network distance measures. To quantify network flow distance, we used the graph theoretical measurement of communicability (Fig. 4). Communicability was chosen over shortest path length (often used in fMRI network analyses) because, in contrast to the shortest path length, it takes into consideration all the possible paths between node pairs. Communicability, as introduced by Estrada & Hatano, can only be used in binary graphs (Estrada & Hatano, 2008). Since each subject’s GC-PCR network is weighted, we used the modified approach described by Crofts & Higham that expands communicability to weighted graphs (Crofts & Higham, 2009). Briefly, this expansion normalizes the weighted graph to diminish the excessive influence of highly connected nodes in the original formulation. This normalization takes place by dividing each weight of the adjacency matrix by the square root of the product of the node strengths of the two nodes that the weight connects. This approach can handle only positive weights, so the weighted graph was thresholded at 0 before calculation. Higher communicability means easier (fewer steps) communication/flow of information/activity between nodes.

**Figure 4:**
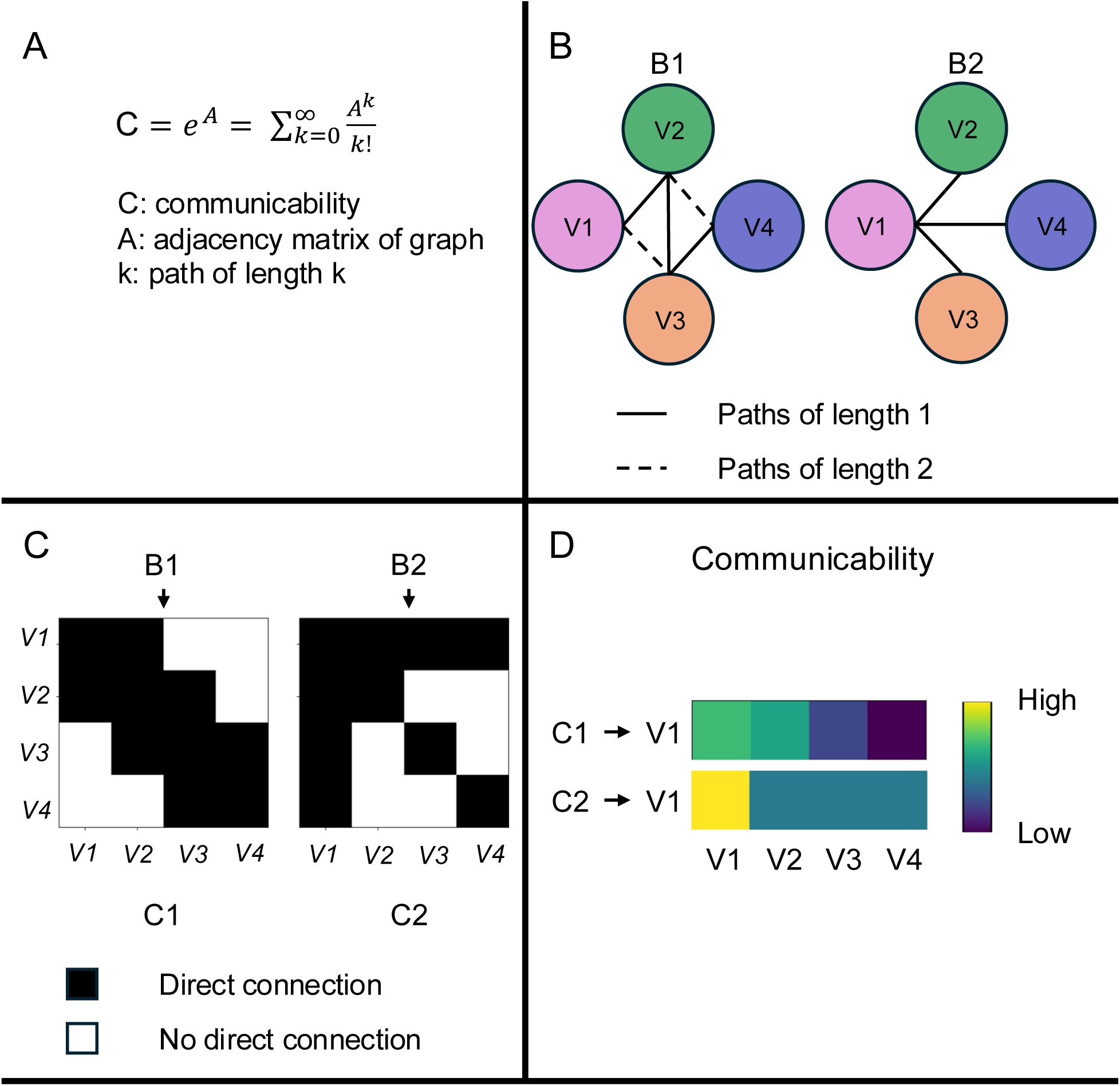
Network communicability captures hierarchical structure via direct and indirect. A: Communicability quantifies the ease of information exchange between nodes in a network by weighting walks of all lengths (Estrada & Hatano, 2008). B: Two toy networks (B1, B2) over V1-V4 illustrating direct (length 1) and indirect paths (length 2). B1 features the hierarchy V1→V2→V3→V4. C: Corresponding adjacency matrices (C1, C2). Black cells indicate direct connections; white indicate no connection. Each row and column are a brain region V1-V4. Here we also assume self-connections. D: Communicability with respect to V1 differs across the two networks, reflecting contributions from both direct and indirect pathways. In graph B1 it recovers the hierarchy with a monotonic decline from V2 to V4. In graph B2, communicability is equal between all nodes with respect to V1, since V1 is connected to all paths.

We defined as network flow distance the communicability of each region of the early visual system with respect to V1. We computed communicability separately for each hemisphere (Fig. 6). We calculated communicability for both parcel-level graphs and vertex level graphs. For the vertex level, we took the mean communicability between all vertices in V1 and all vertices within each target region, resulting in a single number for each region-V1 pair. For statistical comparison between regions, we used a paired t-test, Bonferroni-corrected for multiple comparisons.

**Figure 5:**
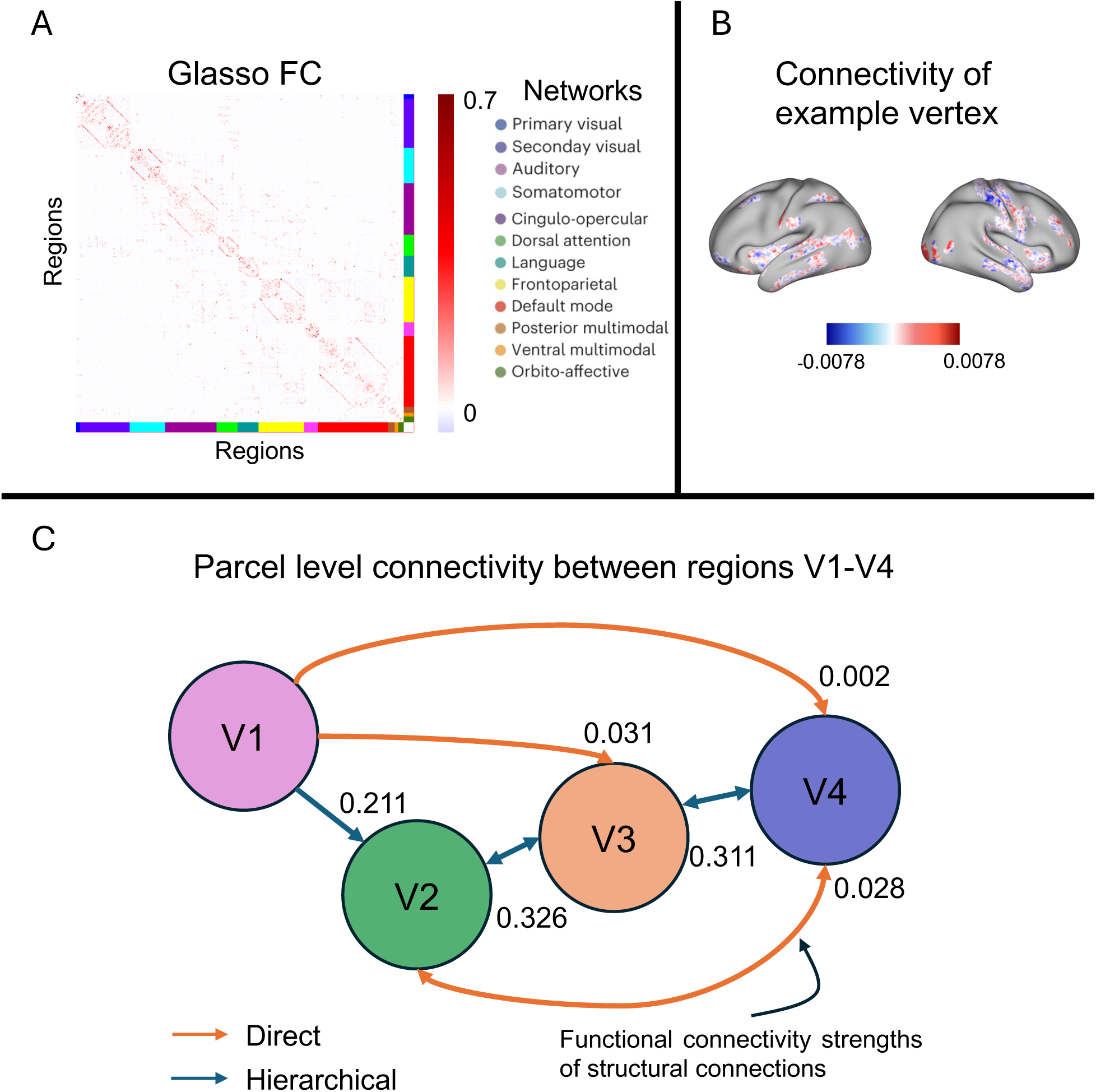
Resting-state functional connectivity at the parcel and vertex levels, focusing on regions V1, V2, V3, and V4. A: Parcel level resting-state FC calculated based on the Glasso algorithm, which reduces confounding during FC estimation relative to the field-standard pairwise Pearson correlation FC measure (Peterson et al., 2025). B: Whole cortex FC of an example V1 vertex calculated based on GC-PCR (see Methods, Fig. 3). C: Force-directed graph based on parcel level connectivity for regions V1-V4. Functional connections are included after masking by group-average structural connectivity (all connections survived). Distances between nodes represent connectivity strength. Shorter distances represent stronger connections. The strongest connection is between V2-V3 and the weakest is between V1-V4. Numbers denote the cross-subject average connectivity strength. Bidirectional arrows denote feedforward and feedback connections that were included in the V1-initiated ENN. Results are based on the right hemisphere. For left hemisphere, see Supplementary Fig. S1. For a full list of percentages see text.

**Figure 6:**
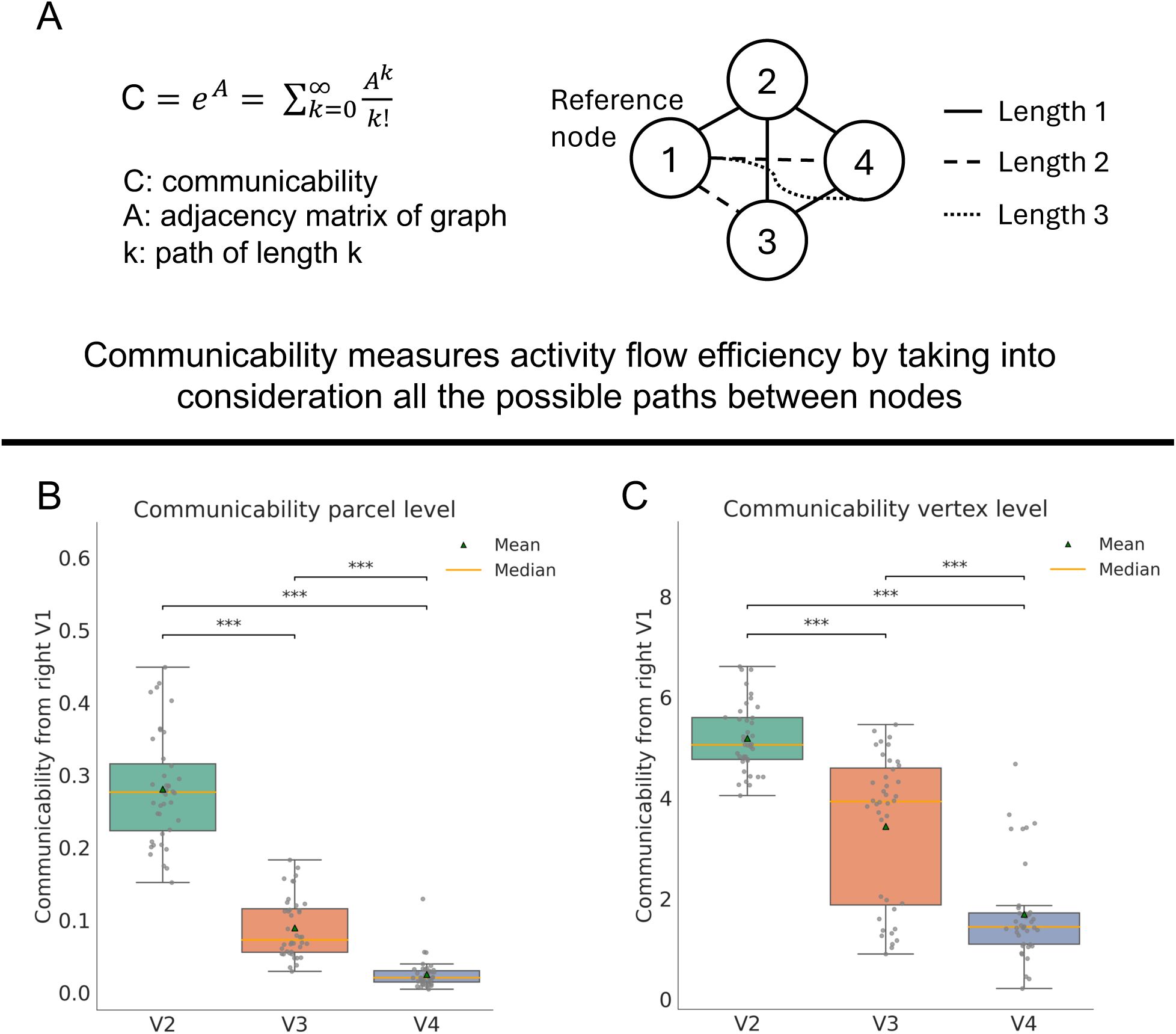
Connectivity-based hierarchy quantified as communicability. A: Definition of communicability and a toy graph illustrating the possible paths between a reference node and the rest of the network. Path of length 2 is 1-2-4 whereas path of length 3 is 1-2-3-4. B: Communicability results based on the parcel level FC graph for the right hemisphere. All pair differences are significant. C: Communicability results based on the vertex level FC graph for the right hemisphere. All pair differences are significant. Values are multiplied by 10^4^ for visualization (vertex FC weights ∼ 10^-4^). Paired t-tests Bonferroni-corrected were used across subjects for all comparisons. Detailed statistics Tables 1,2. For left hemisphere results, see Supplementary Fig. S2. For replication cohort, see Supplementary Fig. S7. Asterisks denote significance: *=p<0.05, **=p<0.01, ***=p<0.001.

### Temporal representational similarity analysis

We extended the standard representational similarity analysis approach developed by Kriegeskorte (Kriegeskorte et al., 2008) to accommodate time points in place of experimental conditions, which we term *temporal representational similarity analysis*. The goal was to avoid the need to define clear boundaries between stimulus categories, better accommodating the continuous nature of the retinotopic mapping used here. This approach was similar to other extension of representational similarity analysis to the temporal domain (Chen et al., 2018; King & Dehaene, 2014).

We calculated cross-validated temporal representational similarity matrices (RSMs) for each of the regions V1-V4 for both the left and right hemisphere. Cross-validation of RSMs has been shown to be important for avoiding biases, such as noise systematically reducing RSM amplitude (Kriegeskorte et al., 2008). We split each of the 6 retinotopic runs into two identical halves based on the stimuli presented. This was possible due to the periodic nature of the stimuli. To achieve the two identical runs, we kept timepoints from the 22^nd^ to the 277^th^ (256 timepoints) for the first four retinotopic runs. Since the last 2 runs were repeated (moving bars), we did not need to split them. To create the cross-validated temporal RSMs, for each subject and for each region, we correlated the first halves across all runs (and the whole 5^th^ run) with the second halves of all the runs (and the whole 6^th^ run) using Pearson’s correlation. This resulted in an 812x812 matrix where each row corresponded to a timepoint from the first half of the runs and each column corresponded to each timepoint of the second half of the runs. Each element of this matrix was the Pearson correlation between the vertices’ activations for a timepoint of the first half of the runs with the activations of another timepoint for the second half of the runs. This approach renders the diagonal of the RSM meaningful since it contains the correlation between the activations at equivalent timepoints between the two halves of the runs (i.e., its repeat reliability). This is especially important for dimensionality measures, which rely on diagonal values in RSMs for accurate estimation.

We additionally calculated a pixel RSM – a temporal RSM based on the similarity of pixel patterns across time points, reflecting the objective spatiotemporal relationships between visual stimulus patterns. We followed an identical approach to calculate the pixel RSM as the brain temporal RSMs. First, we recreated the stimuli used by Benson and colleagues (Benson et al., 2018). We started with the retinotopic stimuli time series of each run. Following (Benson et al., 2018), we superimposed random images of random sizes on the stimuli presented. Since the appearance of those images and their position is random in the original paper, we followed the same approach. The frequency of the presented stimuli was 15Hz, resulting in time series with 4500 timepoints. To bring the timing of the time series to the level of the fMRI signal we down sampled the time series by taking the average signal every 15 timepoints (frequency = 15Hz). This resulted in 300 timepoints for each run. For a detailed description of the methodology and the stimuli, please see (Benson et al., 2018).

In order to create the pixel RSM (Fig. 7A), instead of vertices’ activations we used the pixel activations at each timepoint. Since at each timepoint we had a stimulus of 768x768x3 pixels, where 768 is the width and height of the image and 3 are the color channels, we multiplied for each second those values resulting in a feature vector of 768x768x3 features for each timepoint. This served as the pixel population at each timepoint, equivalent to the vertices’ population. An extra processing step we took for the pixel RSM had to do with rest periods. Since in the timepoints we included, rest periods were present (no stimuli presented), all the pixel values there were constant resulting in Nan values for the correlation. We set correlation values to 1 when both correlated timepoints depicted no stimuli and 0 when one timepoint depicted a stimulus and the other did not. To create the HRF-convolved RSM (Fig. 7B), we convolved the pixel time series of each run described above with a hemodynamic function described in Benson and colleagues. Briefly this HRF was obtained by taking an impulse response that can be computed using spm_hrf[0.1,[6.68 14.66 1.82 3.15 3.08 0.1 48.9]) from SPM. This impulse response was convolved to predict the response to a 1-s stimulus, resampled to a 1-s sampling rate using cubic interpolation and normalized to have a peak amplitude of 1.

**Figure 7:**
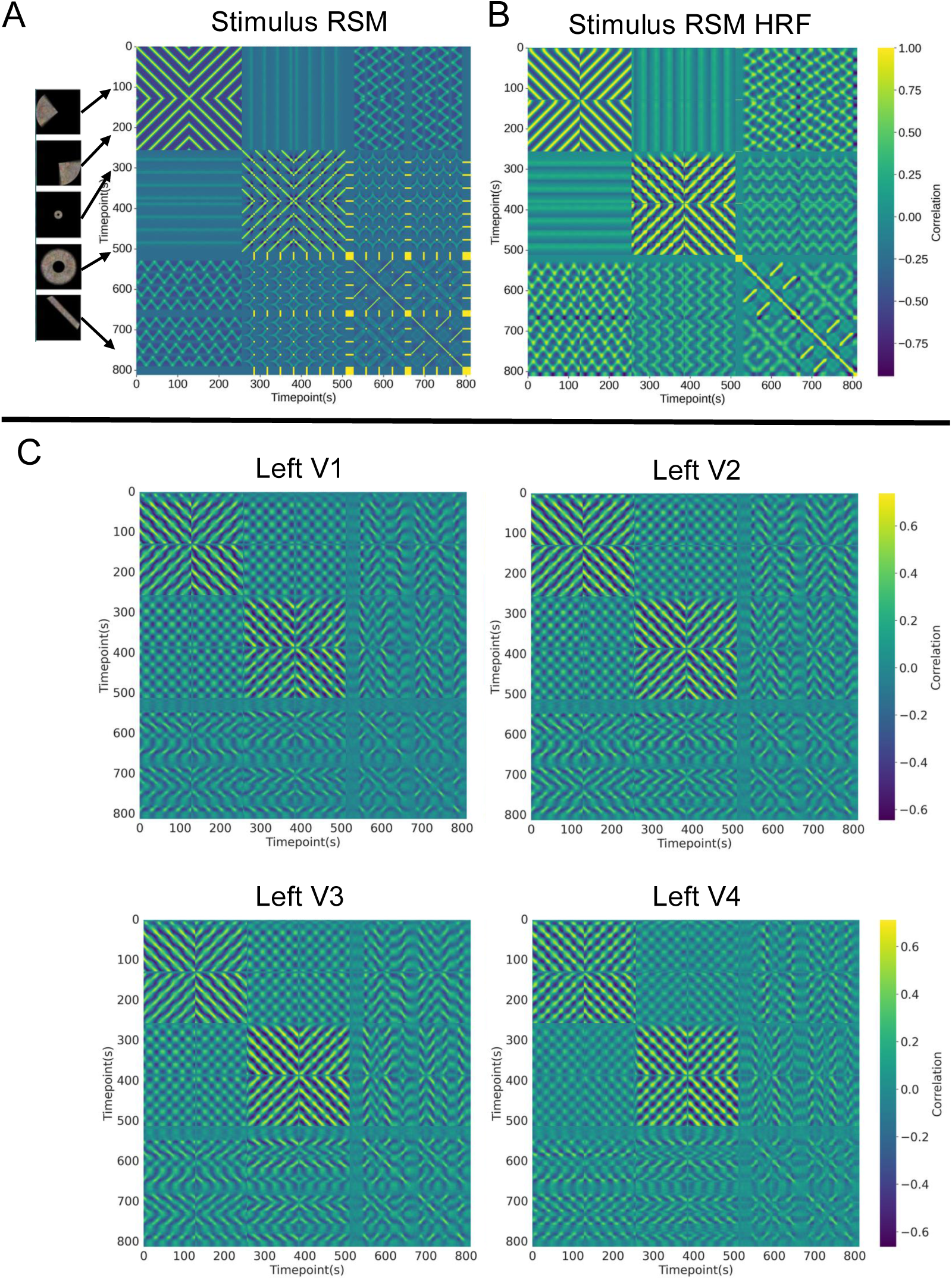
Representational similarity matrices for visual stimulus (pixel) and neural time series. A: RSM based on the pixel time series of the stimuli presented to subjects during the different retinotopic runs. Runs are concatenated. On the left panel example stimuli presented at specific timepoints are depicted. B: Same as A, but pixel time series are convolved with the canonical hemodynamic response function before RSM creation to better match fMRI data (see Methods). C: Cross-validated RSMs for each region of the early visual system for the left hemisphere. For right hemisphere RSMs, see Supplementary Fig. S3.

**Figure 8:**
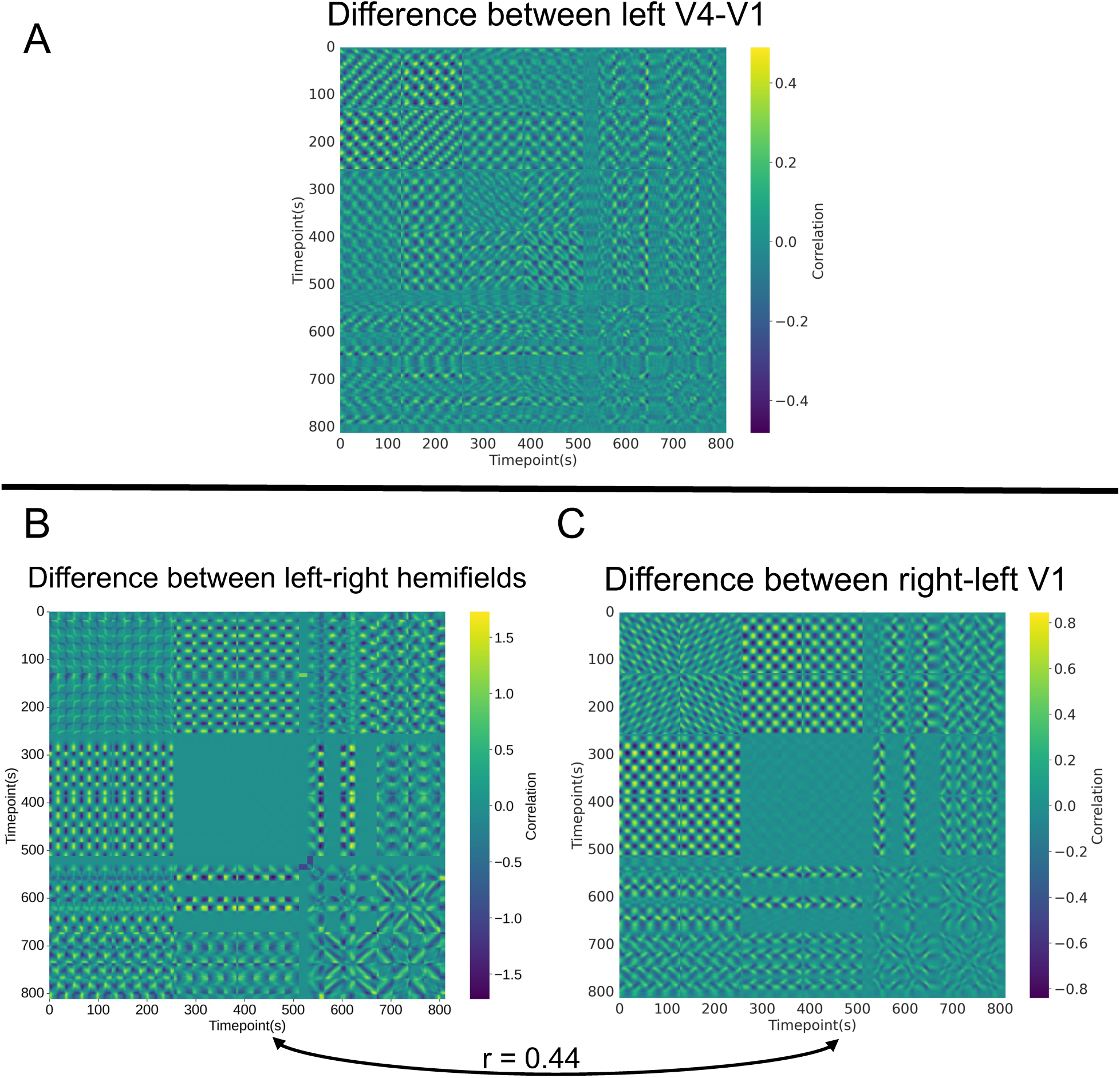
RSM differences reflect underlying cross-region representational differences. A: Differences between left V4 and left V1 representations are present, and the representational difference pattern is reliable across subjects (mean r = 0.195, t = 27.14, p<0.0001). We define representational distance in subsequent analyses based on these different representations. B: Pixel-based RSM demonstrating the difference between visual representations of the left and right hemifields (Methods). The RSM calculated from the time series based on the pixels in the right visual hemifield (right side of the screen) was subtracted from the RSM calculated from the time series based on the pixels in the left visual hemifield (left side of the screen). C: Difference between the representations of left and right V1. Average RSM across subjects. It is evident that this difference in cortical representations captures, especially for runs 3-4 of the retinotopic task (timepoints: 256-512), the difference observed in the pixel RSM. Pearson’s correlation, calculated from the upper triangles (including the diagonal) between the RSMs in B and C, was 0.44 (p<0.0001).

### Representational distance

After calculating for each subject and for each region the temporal RSM, we defined representational distance from V1 as 1 – r (Fig. 9A). r here denotes the correlation between the upper triangles of V1 (separate for each hemisphere) and each other region. Since diagonals of each RSM contain meaningful values due to cross-validation, they were also included in the upper triangles. To assess statistical significance between the distances of regions from V1, these correlations were Fisher z-transformed and paired t-tests were run. The p-values were Bonferroni corrected for multiple comparisons since multiple regions’ pairs distances were compared. Pearson’s correlation was used for all correlations between brain RSMs and between brain and pixel RSMs.

**Figure 9:**
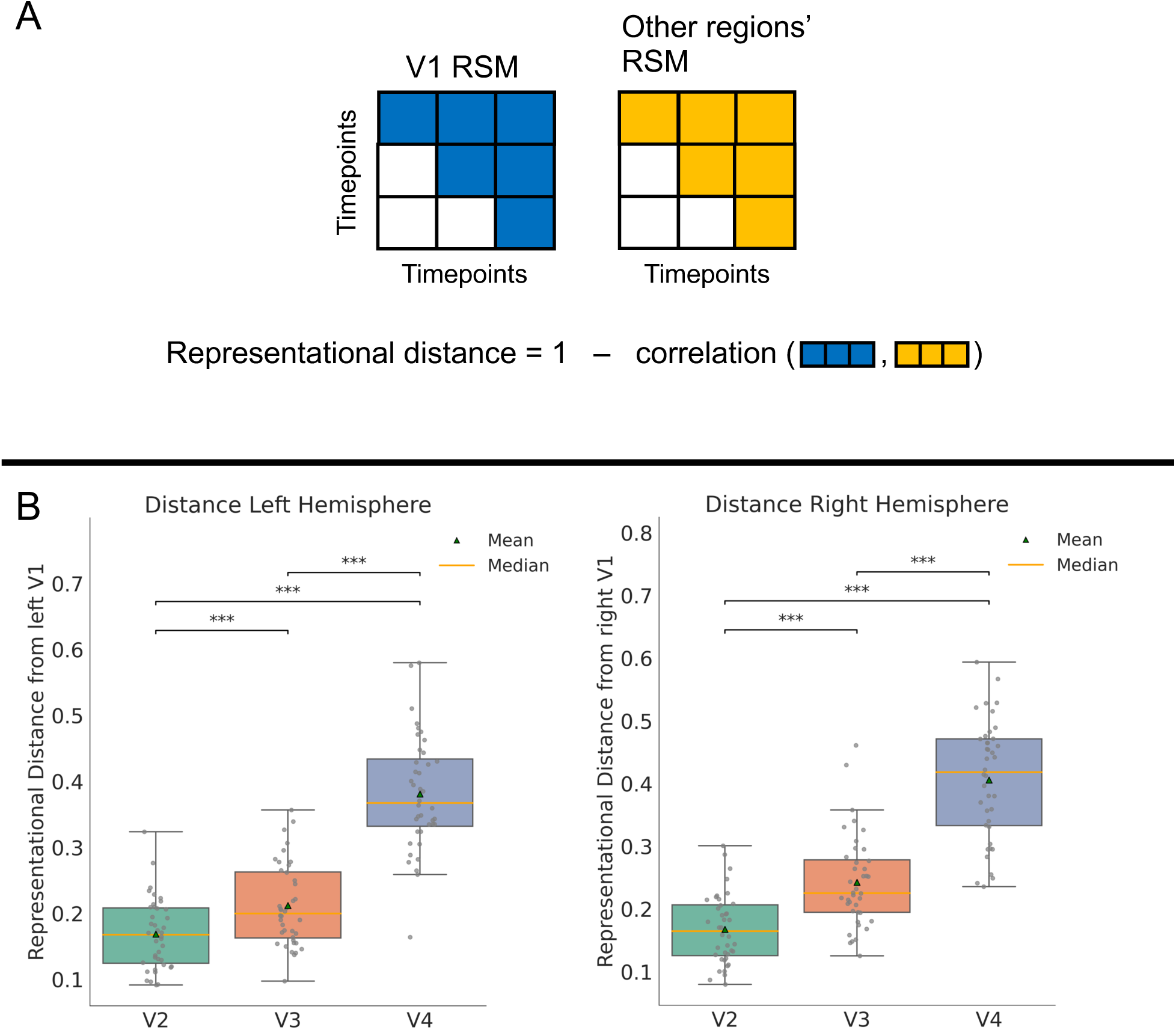
Representational hierarchy based on representational distance from V1. A: Representational distance is defined as the dissimilarity between neural representations (stimulus-evoked activity patterns) of each parcel and V1 for each hemisphere (1-r). B: Representational distance for the left and right hemispheres, respectively. The pattern is similar across both hemispheres, and the same hierarchy emerges: V1→V2→V3→V4. All region pair comparisons are Bonferroni corrected for multiple comparisons (Table 3). For replication cohort, see Supplementary Fig. S7. Asterisks denote significance: *=p<0.05, **=p<0.01, ***=p<0.001.

### Hierarchical and direct contributions

We quantified the contributions of the direct and hierarchical pathways using *in silico* lesions on our V1-initiated activity flow model. For the direct pathways, we lesioned V2-V3 and V3-V4 connections. For the hierarchical pathways, we lesioned V1-V3 and V1-V4 connections. Please note that contralateral connections were also lesioned in both those scenarios (for example left V1 to right V4 as a direct connection). We decided not to lesion V2-V4 connection to maintain the same number of lesions between the two models. For each of the resulting pathways, we ran the V1-initiated activity flow model for 6 steps (matching the number of steps for the original model). We calculated R^2^ between the generated and original time series following a similar approach to the original model. We averaged for each parcel the generated time series and compared them to the original averaged time series of each parcel. Results are shown in figure 12. To statistically compare the R^2^ between the two models at each step, we used a paired Wilcoxon test at each step and then corrected it for multiple comparisons (6 steps) using Bonferroni.

### Dimensionality

We quantified dimensionality as the participation ratio of the RSMs (Chakravarthula et al., 2025; Gao et al., 2017). Participation ratio is given by Equation 3, and it is computed based on the real eigenvalues (λ) following eigen decomposition. We used paired t-tests to compare participation ratios across subjects between regions and a paired Wilcoxon test to compare between models.

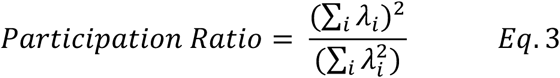

### Hierarchy index

We defined a hierarchy index (HI) for each vertex to quantify the extent that the explained variance of the generated activation for each vertex depends more on the hierarchical or the direct pathways. HI was defined based on Equation 4. HI can range from -1 (full reliance on the direct pathways) to +1 (full reliance on the hierarchical pathways). We then computed Spearman’s correlations between spatial vertex patterns of HI with eccentricity, angle values and receptive field sizes. For eccentricity, we calculated HI based on the third and fourth runs where the stimuli presented are intended to elicit eccentricity contrasts (expanding/contracting rings). We included only those runs to elicit the highest contrasts. For angle, we included the first and second runs. All runs were included for the receptive field size HI. The correlations were Fisher’s z-transformed, and a one-sample, one-sided t-test was run for each region to assess statistical significance. Please note that only regions V3 and V4 (for both hemispheres) were included since there are not separate hierarchical and direct pathways for V2 (V1-V2 connections are both direct and hierarchical). P-values from the t-tests were Bonferroni corrected for multiple comparisons. Angle, as expected, did not show any relationship with the HI.

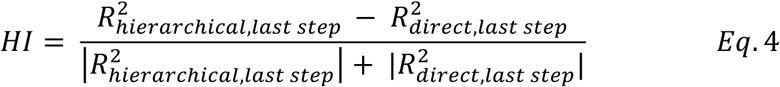

### Difference in deviation of pRF estimates between hierarchical and direct models

As a sanity check, we also tested whether the hierarchical (respectively the direct) pathways are indeed more successful in generating large receptive field sizes and peripheral representations (respectively small and foveal). After generating pRF measurements for the hierarchical and direct models, we calculated the deviation of those measurements from the actual ones focusing on eccentricity and receptive field size. This deviation was calculated at the vertex level for each model as the absolute value of the difference of the log-transformed generated and actual measurements. We applied a logarithmic transformation to diminish the effect of proportional errors. Next the deviation of the hierarchical model was subtracted from the deviation of the direct, resulting in the difference of deviations. Larger difference meant that the generated values from hierarchical model were closer to the observed measurements. We then correlated this difference to the receptive field size and eccentricity using Spearman’s correlation. Equivalent statistical process as the one described above was also followed here.

For statistics we used the scipy stats library from Python. R^2^ was calculated using the sklearn library. For circular correlation we used circcorrcoef from the astropy Python library. All t-tests and Wilcoxon tests were two-sided unless otherwise stated. Degrees of freedom are 39 (40-1) unless otherwise stated. Effect sizes were calculated using standard definitions for one-sample and paired t-tests. For Wilcoxon signed-rank tests, we report signed-rank statistic W and effect size, computed as the normal approximation z statistic divided by the square root of N. N denotes the number of non-zero paired differences.

## Results

The visual system is thought to be hierarchical, but the exact nature of this hierarchy and the role of direct (non-hierarchical) pathways has been a subject of debate. In this study, using fMRI-based retinotopic mapping and connectivity estimates in humans, we tested the role of hierarchical and direct pathways in the generation of visual functionality.

The account of hierarchy we develop here is consistent with many existing definitions, but gains clarity by defining hierarchy in terms of a fundamental, unifying mechanism: activity flows (Figure 1A). Over the past decade, we and others have developed the activity flow framework, which bases the generation of brain functionality (e.g., population receptive fields) on activity flows – the movement of neural activity over network connections (Cole et al., 2016; Ito et al., 2020a; Yan et al., 2021). This framework is both theoretical (gaining clarity through conceptualizing brain functionality in terms of activity flows) and methodological (building models of brain function in terms of activity flows). The activity flow framework has been used to accurately predict task-evoked activity in over 20 studies, involving hundreds of brain regions, dozens of tasks, and multiple neural recording approaches (electroencephalography, multi-unit recording, fMRI, diffusion MRI, and invasive electrocorticography) [see (Cole, 2024) for review].

Connectivity-based and representation-based tests, along with generative activity flow models that combine connectivity and representation data, were used to test our core hypothesis: that hierarchical and direct connectivity patterns in the early visual system generate complementary aspects of visual functionality. This hypothesis was tested via the following specific predictions: (1) that hierarchy in the early visual system can be recovered based on intrinsic connectivity (network flow distance) and also based on retinotopic neural representation changes (representational transformation distance), with the two definitions converging on the classic hierarchy; (2) that both hierarchical (V1 V2↔V3↔V4) and direct (V1↔V3, V1↔V4) V1-initiated pathways contribute to the generation of cortical visual functionality; (3) representations propagated via the hierarchical pathways have lower dimensionality than those propagated via the direct pathways and (4) the hierarchical pathways contribute more than the direct pathways to the generation of large receptive fields and peripheral visual representations. These hypotheses were tested using the novel GC-PCR FC estimation method described above (Fig. 3), graph theory (Fig. 4), multivariate pattern analysis (Fig. 2B), and a biologically plausible V1-initiated ENN (see Methods).

GC-PCR FC yields confound-controlled connectivity estimates at a higher spatial scale than most RSFC studies. This method is based on findings demonstrating that the field-standard pairwise Pearson correlation RSFC approach is strongly confounded by third variables driving false connections (e.g., B←A→ causing false B–C connections). We recently demonstrated that these confounds can be reduced substantially using regularized partial correlation or regularized regression, yielding more direct region-to-region functional connections (Peterson et al., 2025). Notably, the directness of these region-to-region connections were validated using diffusion MRI tractography, which is thought to yield more direct (structural) connections than standard functional connectivity approaches. Our approach used regularized partial correlation (graphical lasso) at the level of brain regions, which then constrained vertex-level RSFC estimation using regularized regression (PC regression). This two-step procedure improved the computational tractability of the confound-controlling procedure, yielding a whole-cortex vertex-wise (dense) functional connectome for each subject. As before, we also validated our connections using DWI tractography, increasing confidence that our direct pathway estimates were not false positives due to network confounding.

The methods used to define hierarchical organization based on graph theoretical flow distance and representational transformation distances from the cortical visual input region are independent, since the first is based on the network architecture as defined by FC (from resting-state fMRI data) whereas the latter is based on the retinotopic (stimulus-evoked) representations (from task fMRI data). As will be seen in the following two sections, the hierarchy defined by both approaches converges to what has been previously suggested: V1↔V2↔V3↔V4. However, direct (non-hierarchical) pathways are also present and strongly contribute to visual processing.

### Network flow distance

We began by testing for the existence of hierarchy in the early visual system based on network architecture via the graph theoretical concept of communicability. Communicability (Estrada & Hatano, 2008) measures how well two nodes in a network can exchange information, considering all possible paths between them, not just the shortest. Information exchange is quantified via simulating all possible walks/flows through the network. By weighting shorter, higher-strength paths more heavily (but still considering all possible paths), it provides a direct index of how “close” (higher communicability) any visual area is to V1 in terms of flow efficiency.

In Figure 5, we illustrate the results from parcel-level connectivity and GC-PCR. In panel C, we see the strengths between the connections for all early visual cortex parcels, delineating all the possible pathways that communicability takes into consideration. It is evident that hierarchical connections are stronger than the direct and, while most subjects have all of these connections, a subset of subjects do not exhibit some direct connections. Specifically, 100% of subjects feature all the hierarchical connections, 80% feature the direct V1-V3 connection, 72.5% the direct V1-V4 and 87.5% V2-V4. These results are based on the right hemisphere, taking into consideration connections originating from either hemisphere (For left hemisphere results, see Supplementary Fig. S1).

We calculated communicability both at the parcel level (based on Glasso-FC; Table 1), and at the vertex level (based on our GC-PCR method; Table 2). Both approaches yielded the same result, with region V2 demonstrating the highest communicability with V1, followed by V3 and finally V4 for both hemispheres (Fig. 6, boxes = median±IQR, gray dots = individual subjects. For left hemisphere results, see Supplementary Fig. S2. For replication cohort, see Supplementary Fig. S7). This corresponds with the field-standard account of hierarchy in early visual cortex (V1↔V2↔V3↔V4).

**Table 1.** Summary of communicability differences’ statistics at the parcel level.

| Region pairs | Difference | t(39) | Cohen's d | P-value |
| --- | --- | --- | --- | --- |
| Left Hemisphere |  |  |  |  |
| V2-V3 | 0.174 | 16.322 | 2.581 | $p < 0.0001$ |
| V2-V4 | 0.220 | 18.763 | 2.967 | $p < 0.0001$ |
| V3-V4 | 0.046 | 9.615 | 1.521 | $p < 0.0001$ |
| Right Hemisphere |  |  |  |  |
| V2-V3 | 0.191 | 14.724 | 2.328 | $p < 0.0001$ |
| V2-V4 | 0.256 | 20.876 | 3.301 | $p < 0.0001$ |
| V3-V4 | 0.064 | 12.557 | 1.985 | $p < 0.0001$ |

Communicability between parcels demonstrated significant differences in all cases. All p-values are Bonferroni-corrected for multiple comparisons.

**Table 2.** Summary of communicability differences statistics at the vertex level.

| Region pairs | Difference ($\times 10^4$) | t(39) | Cohen's d | P-value |
| --- | --- | --- | --- | --- |
| Left Hemisphere |  |  |  |  |
| V2-V3 | 1.907 | 7.745 | 1.225 | $p < 0.0001$ |
| V2-V4 | 2.466 | 11.128 | 1.760 | $p < 0.0001$ |
| V3-V4 | 0.559 | 1.731 | 0.274 | 0.274 |
| Right Hemisphere |  |  |  |  |
| V2-V3 | 1.748 | 6.570 | 1.039 | $p < 0.0001$ |
| V2-V4 | 3.494 | 17.504 | 2.768 | $p < 0.0001$ |
| V3-V4 | 1.746 | 7.308 | 1.155 | $p < 0.0001$ |

Communicability between parcels, calculated at the vertex level demonstrated significant differences in all cases except for the pair: left V3-V4. All p-values are Bonferroni-corrected for multiple comparisons.

**Table 3.** Summary of statistics for representational transformation distance.

| Region pairs | Difference | t(39) | Cohen's d | P-value |
| --- | --- | --- | --- | --- |
| Left Hemisphere |  |  |  |  |
| V2-V3 | 0.043 | 6.555 | 1.036 | $p < 0.0001$ |
| V2-V4 | 0.212 | 25.395 | 4.015 | $p < 0.0001$ |
| V3-V4 | 0.169 | 22.659 | 3.583 | $p < 0.0001$ |
| Right Hemisphere |  |  |  |  |
| V2-V3 | 0.075 | 10.737 | 1.698 | $p < 0.0001$ |
| V2-V4 | 0.238 | 30.438 | 4.813 | $p < 0.0001$ |
| V3-V4 | 0.163 | 20.226 | 3.198 | $p < 0.0001$ |

Representational transformation distance between parcels demonstrated significant difference in all cases. All p-values are Bonferroni-corrected for multiple comparisons.

### Representational transformation distance

We next tested for the existence of hierarchy in the early visual system based on the representational similarities of the retinotopic fMRI time series, which likely reflect the amount of information transformation (computation) implemented with each network flow step from V1. This follows from the activity flow framework’s account of neural processing involving neural activity vector (i.e., representational) transformations over multiple steps (Ito et al., 2022). Distance in this case is defined as the dissimilarity (1-r) between the upper triangle vectors (including the diagonal) of each region’s RSM with V1’s RSM (see Methods). To validate that the visual region RSMs are indeed representations of the presented visual stimuli, we first constructed a stimulus RSM by estimating the correlations among the raw stimulus pixel-intensity vectors across timepoints for all retinotopic runs (Fig. 7 A-B). All eight regions’ RSMs correlated significantly with the HRF-convolved stimulus RSM (mean r = 0.3263, all p < 0.0001). This demonstrates that the fMRI data successfully captured meaningful representational structure induced by the retinotopic mapping stimuli.

In Figure 7C we show each brain region’s RSM, highlighting how neural representations transform the raw stimulus similarity pattern (For right hemisphere results, see Supplementary Fig. S3). In Figure 8A we show differences in these representational patterns between V4 and V1. These differences were calculated as the pairwise numerical difference of a pair of RSMs. This difference demonstrates that systematic changes in representation patterns are present across regions, despite their overall similarity with the stimulus pixel patterns.

To test the stability of this difference across subjects, we calculated a mean difference RSM using a leave-one-subject-out approach. For each subject we correlated its RSM difference to the mean difference and ran a one-sample t-test against 0 to test for significance (mean r = 0.441, t = 23.81, p < 0.0001).

Also in Figure 8, we depict the difference in stimulus patterns between left and right visual hemifields (Fig. 8-B, Methods), along with the difference between the representations of left and right V1 (Fig 8-C). The Pearson correlation between the pixel RSM difference pattern and V1 hemisphere RSM difference pattern (r=0.44, t = 35.45, p<0.0001) was statistically significant across subjects, demonstrating that the RSM-estimated neural representations capture veritable differences in visual stimulus patterns.

Representational transformation distances mirrored the connectivity-based hierarchy: mean distance from V1 was lowest for V2, higher for V3 and highest for V4 (Table 3). The distribution of the distance values across subjects for each parcel is displayed in Figure 9 (boxes = median±IQR, gray dots = individual subjects).

### ENN model validation: Modeling flows over empirical connections accurately generates retinotopic time series

Activity flow modeling has been shown to be effective in generating task-evoked activations across a variety of brain regions and tasks (Cole et al., 2016; Ito et al., 2022). This framework treats the brain’s FC as a wired network through which signals move (“flow”) to produce task responses (Ito et al., 2017). This places brain flows at the center of neural computation in general, with a focus here on the generation of visual representations in early visual cortex. To increase the biological plausibility of our ENN model, we based its construction on Cocuzza et al. (2024), allowing only stimulus-evoked activity in V1 to generate activations in regions V2, V3, and V4 (V1-initiated), which increases the validity of directional inferences. Also, our vertex-wise model increases the spatial resolution of flow-based inferences relative to most earlier studies.

While all results were generated at the vertex level, we also report results at the parcel level by averaging across vertices within each parcel. We expected predictions at the parcel level to be more robust compared to the vertex level due to greater repeat reliability of the to-be-predicted data due to cross-vertex averaging (which increases signal-to-noise). Briefly, after generating time series for each vertex, we averaged them across vertices and compared them to the actual averaged time series for each parcel for each run. We then averaged those values across runs. The number of steps was determined as the step wherein the increase of R^2^ asymptotes (see Methods). At the 6th step group-averaged R^2^ plateaued, suggesting the system had reached a steady state (Fig. 4).

The generated time series showed a significant correspondence with the actual time series (Table 4; Fig. 10. For replication cohort, see Supplementary Fig. S8 and Supplementary Table S1). Generated-to-actual similarities were significantly above chance for all six regions (all p<0.0001). The highest correlation at the last step was observed for right V2 (r = 0.85), followed by left V2 (r = 0.84). The lowest correlation was observed for right and left V4 (r = 0.65 and r = 0.68, respectively). Example plots for a random subject for the second run across parcels are shown in Figure 10 (B-D). R^2^ followed the same pattern as the correlation values.

**Figure 10:**
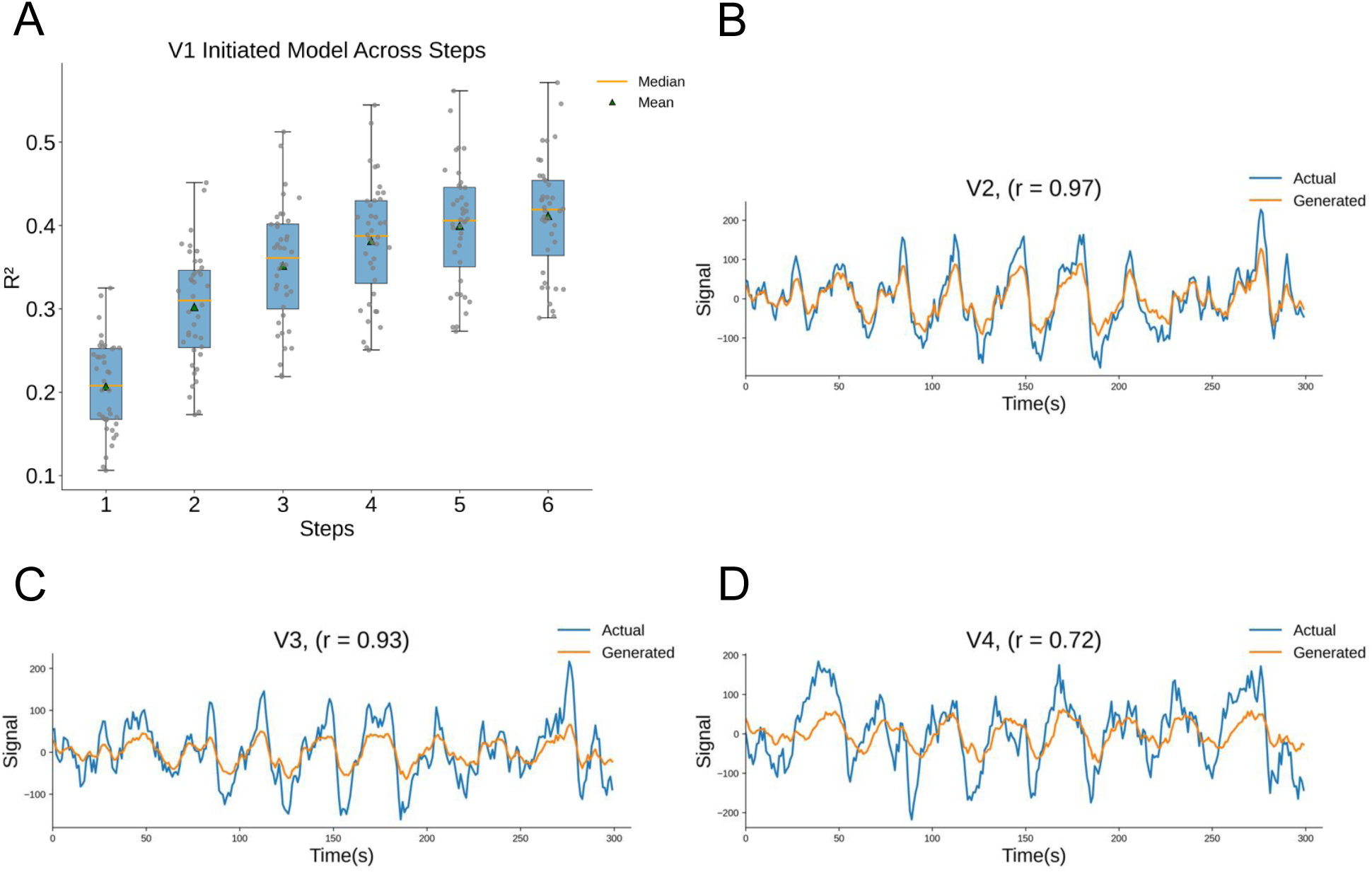
Simulated activity flow using individual-subject FC-based Empirical Neural Networks (ENNs) accurately generates retinotopic time series. A: R^2^ (coefficient of determination) between predicted and actual time series (across left and right V1, V2, V3, and V4) at each flow step of the model. Calculated after averaging generated and original time series across vertices for each parcel. R^2^ was then averaged across subjects and retinotopic runs. Performance plateaus at the 6^th^ step (average R^2^ = 41.134%). B-D: Correlation between the original and the generated time series for a random subject for the second retinotopic run. Target regions of the left hemisphere are depicted. Generated and original vertex time series are averaged for each parcel. For replication cohort, see Supplementary Fig. S8.

**Table 4.** Activity flow model retinotopic time series prediction accuracy (parcel level)

| Region | Pearson's r | $R^2$ (%) | P-value |
| --- | --- | --- | --- |
| Left V2 | 0.844 | 56.29 | $p < 0.0001$ |
| Left V3 | 0.804 | 45.22 | $p < 0.0001$ |
| Left V4 | 0.679 | 26.88 | $p < 0.0001$ |
| Right V2 | 0.852 | 54.65 | $p < 0.0001$ |
| Right V3 | 0.811 | 43.66 | $p < 0.0001$ |
| Right V4 | 0.648 | 20.11 | $p < 0.0001$ |

Pearson $r$ and $R^2$ are calculated across the average time series for each parcel and for each subject. Here we present the average across subjects at the last step of the model. P-values were obtained after running a one-sample one-sided Wilcoxon test on the $R^2$ distribution across subjects. Bonferroni corrected for multiple comparisons.

### ENN model validation: Modeling flow over empirical connections accurately generates pRF parameters at the parcel level

We hypothesized that stimulus-evoked activations flowing over intrinsic FC are sufficient to generate visual functionality in the early visual system. To test this, after generating retinotopic task time series for each vertex, we fed those predictions into a pRF model to generate retinotopic maps at the vertex level, focusing on angle and eccentricity (see Methods). We calculated the circular correlation for angles and the linear correlation for eccentricities between predicted and actual values across subjects at the parcel level, testing for results in each region of the left and right early visual system (Figure 11, boxes = median±IQR, gray dots = individual subjects. For replication cohort, see Supplementary Fig. S9). All region-level correlations were statistically significant (p < 0.0001), except for the angle comparison for right V4 (Table 5). The lowest correlations for eccentricity were observed in right V2 (r=0.354), while the lowest for angle were in right V4 (r=0.120). Conversely, left V4 exhibited the highest correlations, with a mean angle correlation of 0.397 and a mean eccentricity correlation of 0.718. These results further validate the generative accuracy of the activity flow models for retinotopic visual functionality.

**Figure 11:**
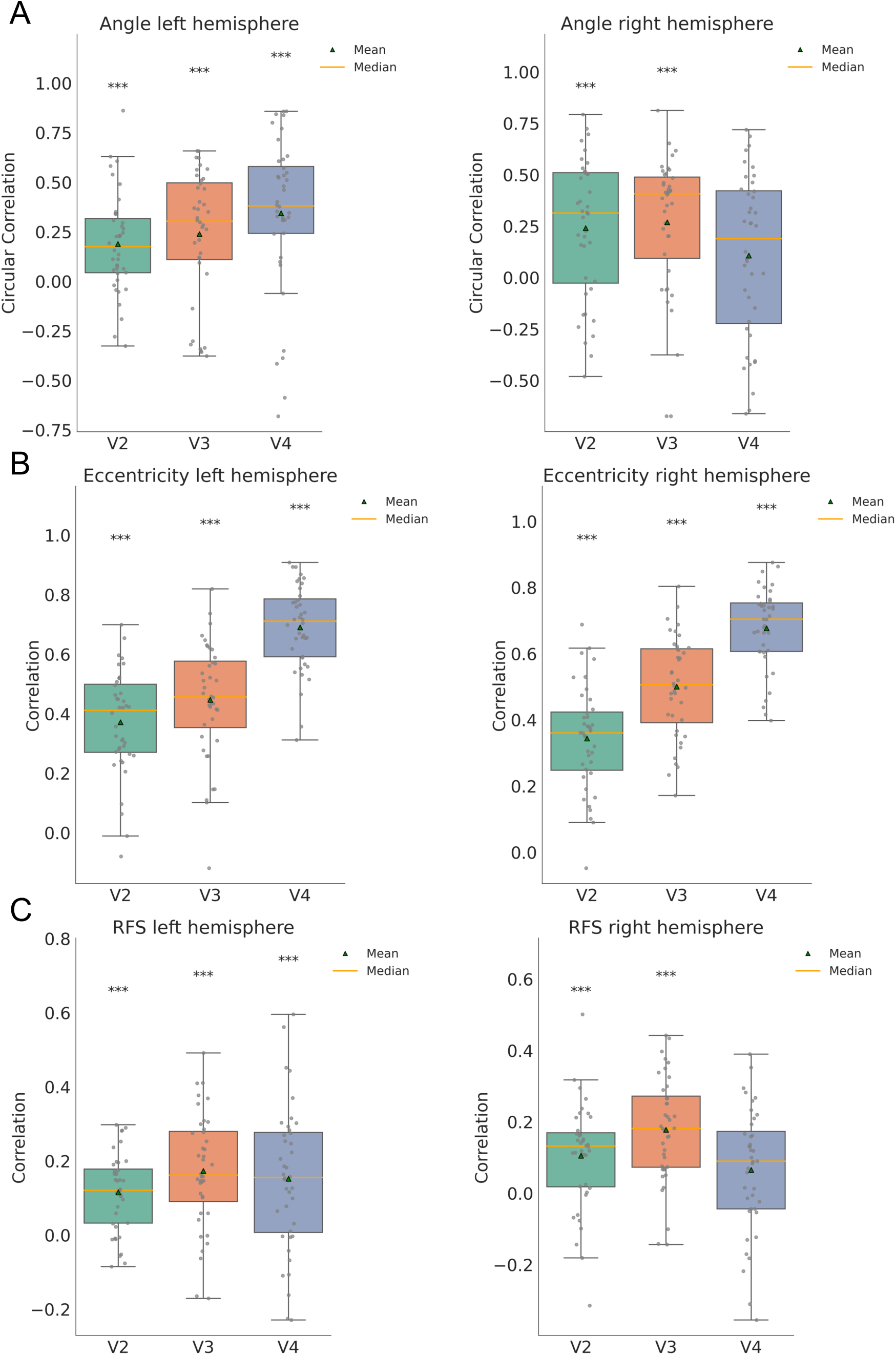
ENN-based simulated flows over FC accurately generate population receptive field (pRF) measurements. A: Circular correlation between actual and generated angle across vertices for each parcel of the early visual system for left and right hemisphere respectively. Correlations are significant for all parcels (p < 0.0001) except for right V4 (Table 5). B: Correlations between actual and generated eccentricity for each parcel of the early visual system for left and right hemisphere respectively. Correlations are significant for all parcels (p < 0.0001, Table 5). C: Correlations between actual and generated receptive field size for each parcel of the early visual system for left and right hemisphere respectively. Correlations are significant for all parcels (p < 0.00041) except for right V4 (Table 5). Boxes = median±IQR, gray dots = individual subjects. For replication cohort, see Supplementary Fig. S9. Asterisks denote significance: *=p<0.05, **=p<0.01, ***=p<0.001.

**Figure 12:**
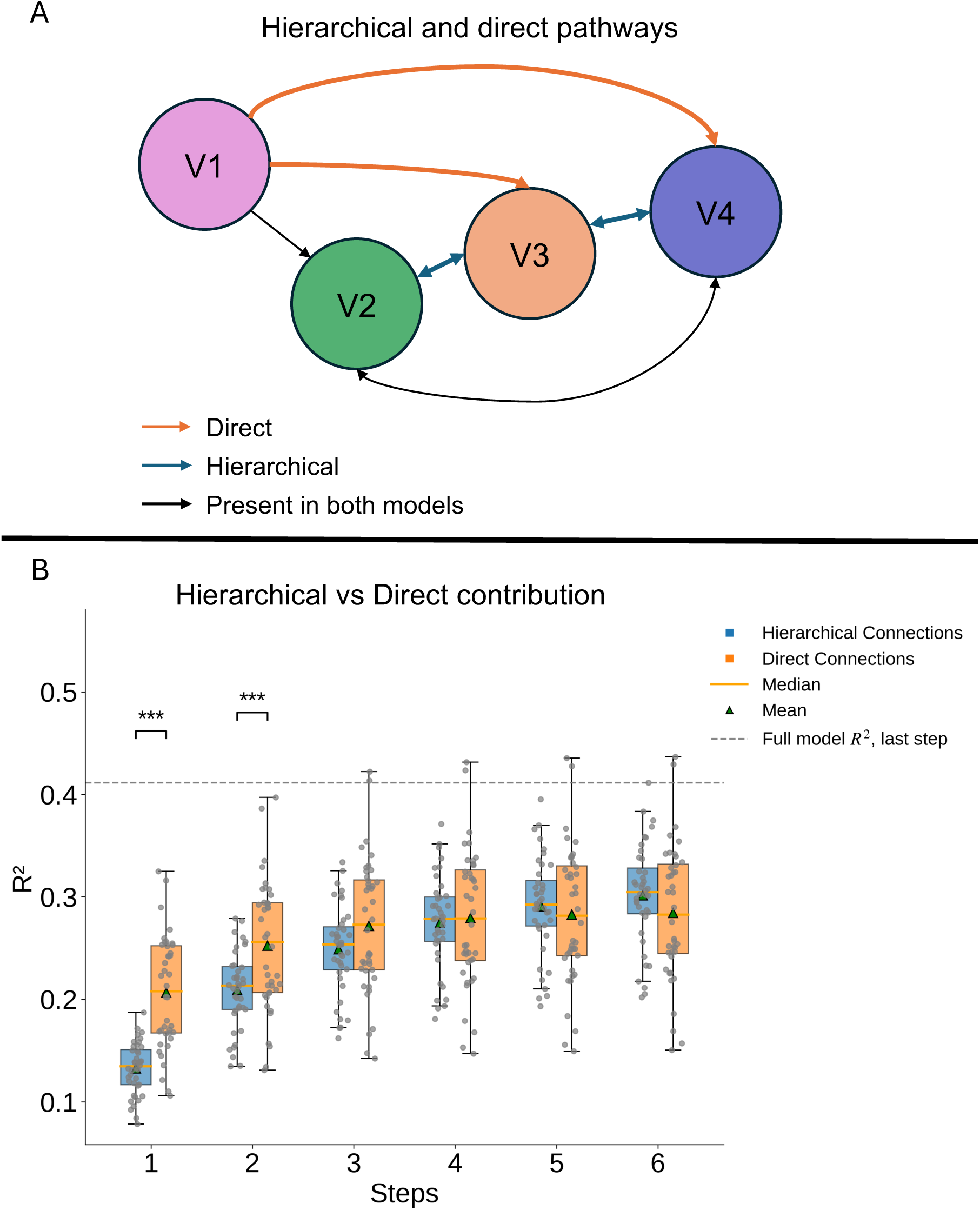
*In silico* ENN lesioning experiment: Both hierarchical and direct connections contribute to visual functionality generation, with direct pathways generating more representational details faster (in fewer steps). A: Modification of the force-directed graph from figure 5-C. Functional connections are included following masking by group-average structural connectivity (all connections survived). Edge length represents connectivity weight between parcels. Larger length denotes weaker connectivity. Hierarchical and direct pathways considered and lesioned in the *in silico* experiment are also shown. For the hierarchical (respectively direct) model, direct (respectively hierarchical) connections are lesioned. B: Comparison between the two different lesion models at the vertex level. Generated time series were averaged for each parcel and then compared to the original ones resulting in R^2^ values. These values were averaged across runs and parcels. Initially, the two models are significantly different with the hierarchical pathways demonstrating larger increase across steps. This shows that as the system reaches its steady state, the hierarchical pathways become an equal contributor to visual functionality generation outperforming the direct ones at the last two steps (difference not significant). Comparisons are computed at the parcel level Bonferroni-corrected (Table 7). For replication cohort, see Supplementary Fig. S10. Asterisks denote significance: *=p<0.05, **=p<0.01, ***=p<0.001.

**Table 5.** Summary of correlations between model-generated and actual values for angle, eccentricity and receptive field size (RFS).

| Region | Angle | P-value | Eccentricity | P-value | RFS | P-value |
| --- | --- | --- | --- | --- | --- | --- |
| Left V2 | 0.209 | $p < 0.0001$ | 0.384 | $p < 0.0001$ | 0.117 | $p < 0.0001$ |
| Left V3 | 0.259 | $p < 0.0001$ | 0.467 | $p < 0.0001$ | 0.177 | $p < 0.0001$ |
| Left V4 | 0.397 | $p < 0.0001$ | 0.718 | $p < 0.0001$ | 0.159 | $p = 0.0001$ |
| Right V2 | 0.270 | $p < 0.0001$ | 0.354 | $p < 0.0001$ | 0.108 | $p = 0.0004$ |
| Right V3 | 0.289 | $p < 0.0001$ | 0.517 | $p < 0.0001$ | 0.182 | $p < 0.0001$ |
| Right V4 | 0.120 | $p = 0.585$ | 0.696 | $p < 0.0001$ | 0.067 | $p = 0.132$ |

Circular and Spearman's correlations are calculated between actual and generated angle, receptive field size and eccentricity respectively. Correlations are calculated across vertices for each parcel. P-values are obtained from one-sample t-tests, Bonferroni corrected for multiple comparisons.

### ENN model validation: Modeling flow over empirical connections accurately generates retinotopic time series at the vertex level

In order to provide a more detailed account of activity flow model performance, we next focused on vertex level model accuracy. In contrast to the parcel-level results, we calculated Pearson r and R^2^ based on model-generated vertex-wise activity patterns. We ran a MaxT approach (see Methods) to correct for multiple comparisons at the vertex level. In the context of these analyses involving many comparisons, this permutation test-based approach is an improvement over Bonferroni correction for multiple comparisons used in other analyses, since (unlike Bonferroni) it is not prone to an excessive type II error. Out of the 2565 vertices of left and right V2-V4, 45.575% of vertices were significant (p<0.05). 42.651% showed p < 0.01 and 40.039% showed p < 0.001.

We also averaged those R^2^ values across vertices for each parcel and ran the MaxT approach at the parcel level. As expected, due to increased noise at the vertex level, model performance was decreased (compared to the parcel level described above) but still significant (p<0.001) for all parcels. The average correlation across parcels was 0.488 (Table 6. For replication cohort, see Supplementary Table S2).

**Table 6.** Summary of Pearson’s r and R^2^ results at the last step of the model.

| Region | Pearson's $r$ | $R^2$ | P-value |
| --- | --- | --- | --- |
| Left V2 | 0.518 | 0.197 | $p < 0.001$ |
| Left V3 | 0.529 | 0.190 | $p < 0.001$ |
| Left V4 | 0.460 | 0.141 | $p < 0.001$ |
| Right V2 | 0.499 | 0.176 | $p < 0.001$ |
| Right V3 | 0.520 | 0.180 | $p < 0.001$ |
| Right V4 | 0.392 | 0.098 | $p < 0.001$ |

Pearson $r$ and $R^2$ are calculated for each vertex for each subject. Here the average $r$ and $R^2$ across vertices for each region and across subjects is reported. P-values based on $R^2$ are obtained from a MaxT nonparametric permutation test.

**Table 7.** Summary of statistics for lesion models’ contributions across steps.

| Steps | Hierarchical Pathways ($R^2$) | Direct Pathways ($R^2$) | p-value |
| --- | --- | --- | --- |
| 1 | 0.132 | 0.207 | $p < 0.0001$ |
| 2 | 0.201 | 0.252 | $p < 0.0001$ |
| 3 | 0.249 | 0.272 | $p = 0.073$ |
| 4 | 0.275 | 0.279 | $p = 0.6656$ |
| 5 | 0.291 | 0.283 | $p = 0.5448$ |
| 6 | 0.301 | 0.284 | $p = 0.1316$ |

Average $R^2$ values for each lesion model across steps. Generated time series were averaged for each parcel and then compared to the original ones resulting in $R^2$ values. These values were averaged across parcels and runs. P-values are obtained from paired Wilcoxon tests across subjects for each step, Bonferroni corrected.

### *In silico* ENN lesion experiments: Hierarchical and direct pathways generate visual functionality collaboratively, with more rapid processing via direct pathways

Based on the hierarchy established by connectivity and representational transformation distances above, we hypothesized that both hierarchical and direct pathways contribute to visual functionality generation. To test this, we lesioned the original V1-initiated activity flow model in two ways: 1) hierarchical lesions (V2↔V3, V3↔V4 connections removed), allowing only direct connections and 2) direct lesions (V1↔V3, V1↔V4 connections removed), allowing only hierarchical connections (see Methods). Note that V2↔V4 was kept in both models to keep the total number of lesions matched between the two lesion models (For lesion analysis where V2↔V4 was removed from the hierarchical model and V1↔V2 was removed from the direct model, see Supplementary Fig. S6). For each model and each propagation step, we calculated R^2^ between the model’s predicted time series and the observed fMRI time series in each target region at the vertex level.

Figure 12 illustrates, for steps 1–6, the distribution of these R^2^ values across subjects (boxes = median±IQR, gray dots = individual subjects. For replication cohort, see Supplementary Fig. S10) for V2, V3, and V4 under each lesion condition. At each step, paired Wilcoxon tests (Bonferroni-corrected for multiple comparisons) were computed between the R^2^ distributions of the two models, demonstrating significant differences for the first 2 steps. Given that each flow step must take time (e.g., due to propagation of action potentials underlying each parcel-to-parcel flow), this suggests the direct pathways were faster than the hierarchical pathways in generating representational details throughout the early visual system.

At the 3^rd^ step, the accuracy of the hierarchical model had reached the level of the direct model, even surpassing it at the 5^th^ and 6^th^ step (though the models were not significantly distinct at these steps) (Table 7). Together, these results demonstrate that both hierarchical and direct pathways are strong contributors to visual information propagation—direct connections drive early-step propagation, while hierarchical connections become increasingly important at later steps as the system reaches its steady state.

### *In silico* ENN lesion experiment: Hierarchical pathways compress visual representations

Prior work has shown that V1 neurons encode high-dimensional simple features (e.g., orientations) and higher-order areas like V4 encode lower-dimensional visual information (e.g., shapes), suggesting visual information compression occurs along the visual hierarchy. Based on our hypothesis that flow steps implement neural computations (e.g., compression), along with fewer flow steps being present in the direct pathways, we hypothesized that the direct pathways would preserve higher dimensional representations compared to the hierarchical pathways. We quantified dimensionality as the participation ratio of each region’s RSM [see Methods; (Chakravarthula et al., 2025; Gao et al., 2017)] and compared across regions: dimensionality decreased significantly from V1 to V4 (Fig. 13; boxes = median±IQR, gray dots = individual subjects; V1 > V4, p = 0.0013, Bonferroni-corrected; Table 8).

**Figure 13:**
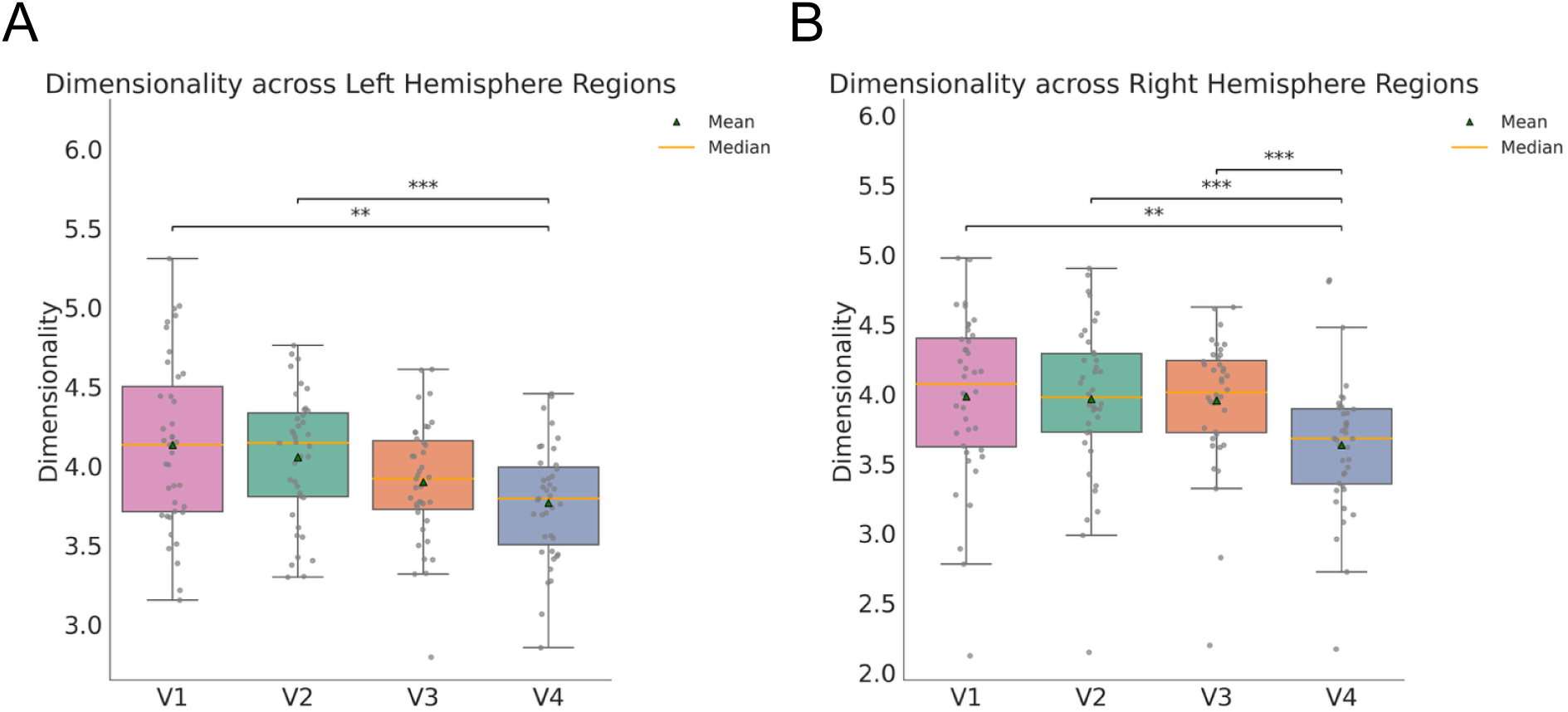
Visual representation dimensionality decreases along the visual hierarchy. A: Dimensionality across parcels for the left hemisphere, independent of the ENN model. Dimensionality is calculated as the participation ratio of each parcel’s RSM. As visual information becomes increasingly less complex (V1→V4), dimensionality decreases. For visualization purposes, we removed an outlier with dimensionality = [1.852, 2.255, 2.926, 1.092] for regions V1, V2, V3 and V4 respectively. B: Dimensionality across parcels for the right hemisphere, independent of the ENN model. Similar trend is observed here. Interestingly, for the right hemisphere V4 demonstrated significant differences with all the other regions (Table 8). P-values are calculated based on paired t-tests across subjects, Bonferroni corrected. Asterisks denote significance: *=p<0.05, **=p<0.01, ***=p<0.001.

**Table 8.**
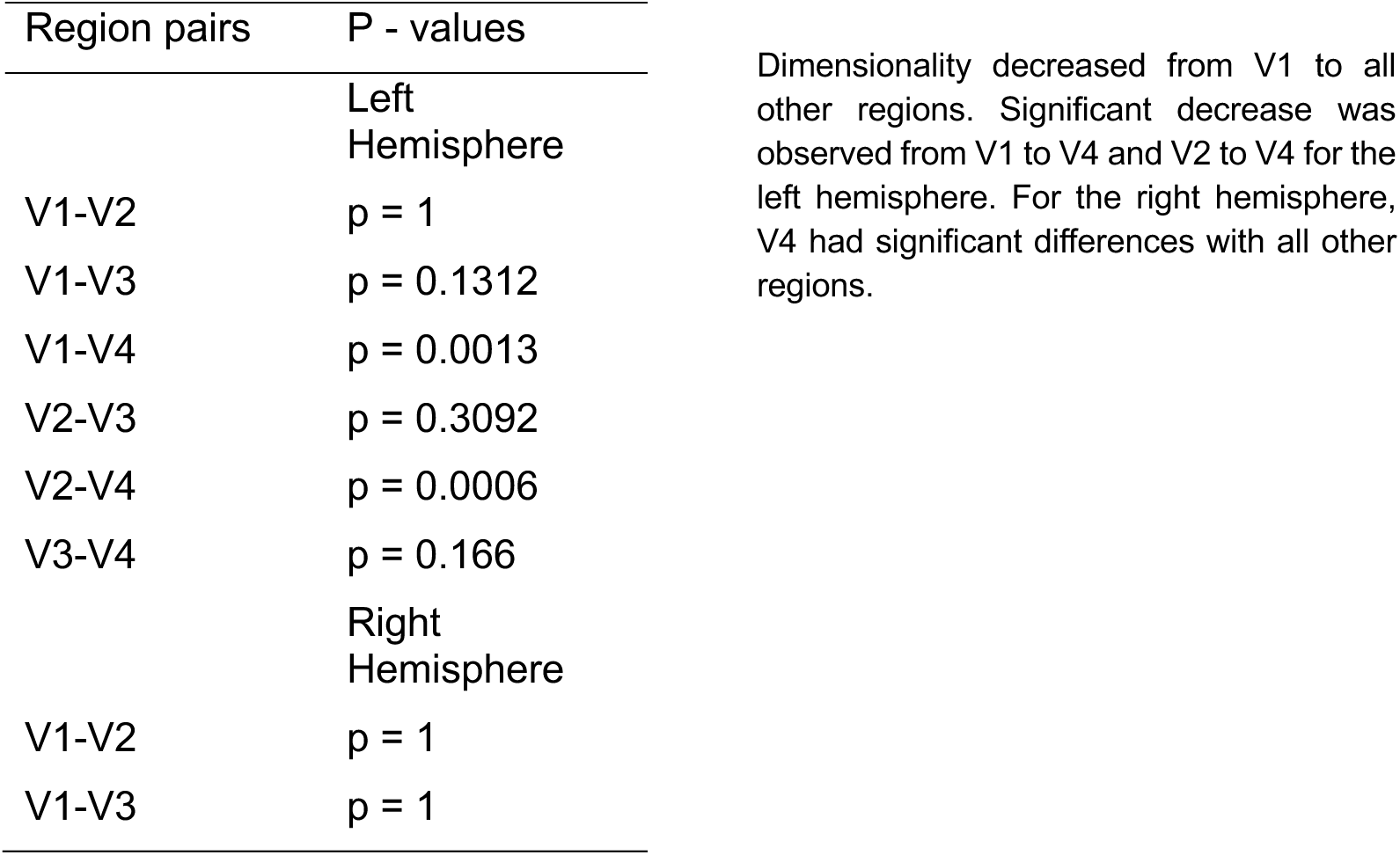
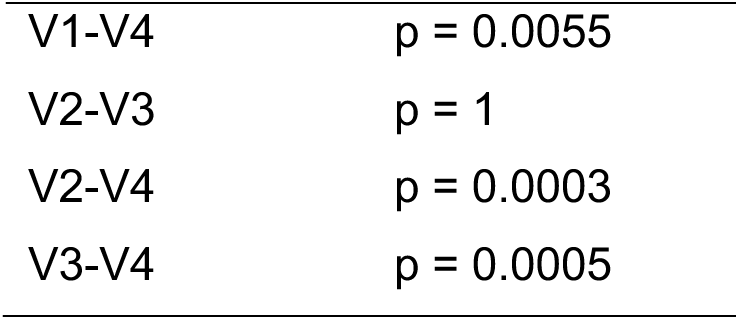
Summary of statistics for dimensionality difference across regions.

Next, we computed dimensionality at each step for both lesioned ENN models (Fig. 14; boxes = median±IQR, gray dots = individual subjects). As hypothesized, the direct pathways retained higher dimensionality across steps, whereas the hierarchical pathways demonstrated decreases as the system reached its steady state (paired Wilcoxon tests on adjacent steps, Bonferroni-corrected. All adjacent comparisons were significant at p < 0.05, except for the second to third step transition). Paired Wilcoxon tests confirmed significant model differences across all steps following the initial step (Table 9).

**Figure 14:**
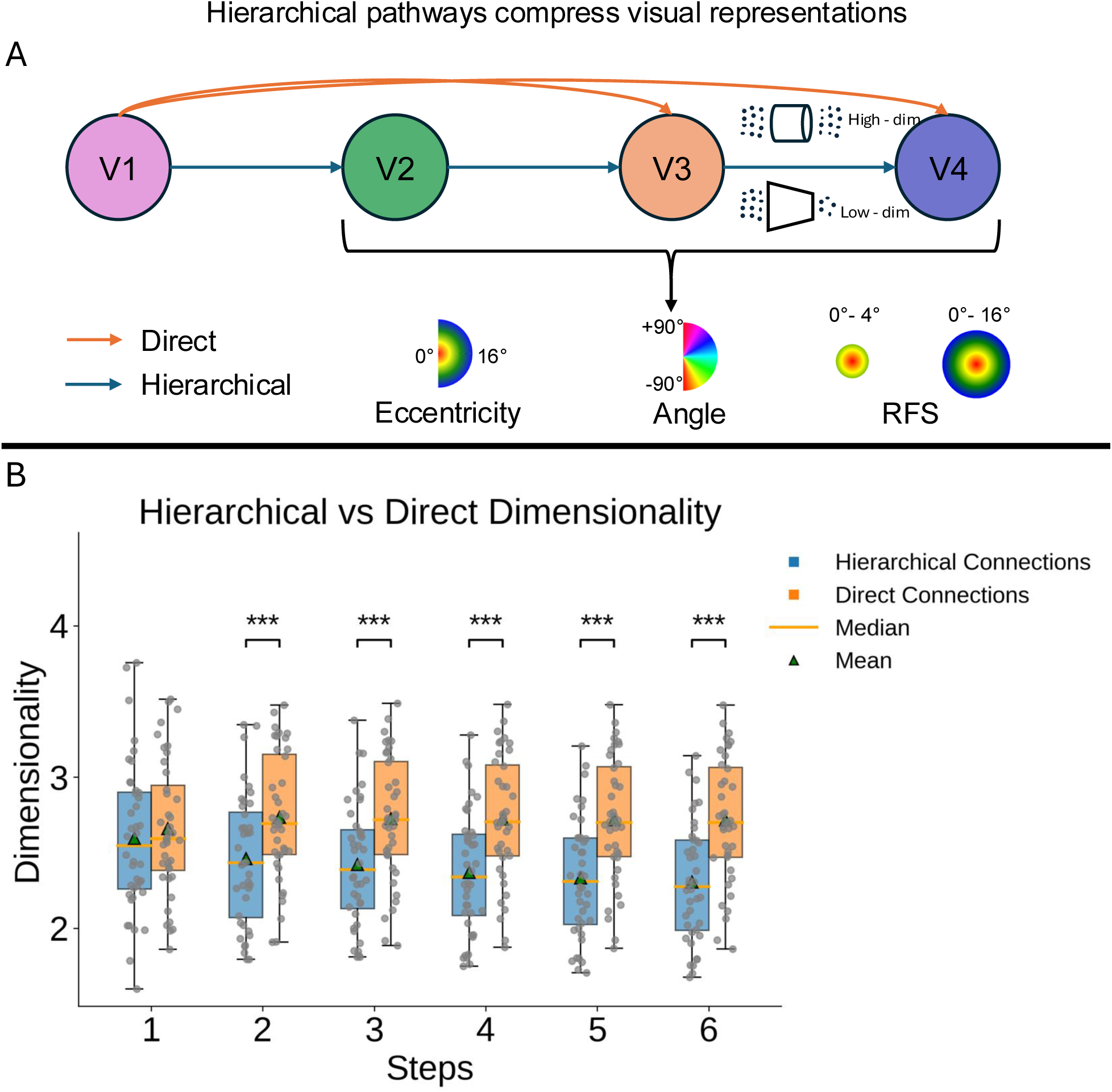
Hierarchical pathways compress representational dimensionality over flow steps, while direct pathways retain higher dimensionality. A: We tested the hypothesis that the hierarchical pathways would compress visual representations. B: Dimensionality across steps for the hierarchical and direct pathways. Excluding the first step, all paired Wilcoxon tests across subjects (Bonferroni-corrected) showed increased dimensionality for the direct pathways. This validates our hypothesis that the direct pathways retain high-dimensional representations from the primary visual cortex. For replication cohort, see Supplementary Fig. S10. Asterisks denote significance: *=p<0.05, **=p<0.01, ***=p<0.001.

**Table 9.** Summary of statistics for dimensionality difference across steps between lesion models.

| Steps | Difference | W | Effect size | P - values |
| --- | --- | --- | --- | --- |
| 1 | 0.064 | 341 | 0.147 | $p = 1$ |
| 2 | 0.274 | 73 | 0.716 | $p < 0.0001$ |
| 3 | 0.302 | 44 | 0.778 | $p < 0.0001$ |
| 4 | 0.353 | 24 | 0.820 | $p < 0.0001$ |
| 5 | 0.384 | 15 | 0.839 | $p < 0.0001$ |
| 6 | 0.411 | 8 | 0.854 | $p < 0.0001$ |

Direct pathways retained higher dimensionality across steps, whereas the hierarchical ones demonstrated slight decrease. P-values are obtained from paired Wilcoxon signed-rank tests across subjects for each step, Bonferroni corrected.

### *In silico* ENN lesion experiment: Large receptive fields and peripheral representations rely more on the hierarchical pathways

It has been previously demonstrated that increased receptive field sizes within the visual system tend to incorporate peripheral representations (Dumoulin & Wandell, 2008). This likely reflects hierarchical compression of visual information into representations integrated across space. Based on this, we next tested our final hypothesis that larger receptive field sizes and more peripheral representations rely more on the hierarchical pathways. To this end, first, we created a hierarchy index (see Methods), which we then tested for correlations with eccentricity and receptive field size for each vertex of V3 and V4 for both hemispheres. The hierarchy index quantifies, for each vertex, the extent to which visual functionality is generated based on the hierarchical pathways compared to the direct pathways.

Consistent with our hypothesis, for regions V3 and V4, all correlations between the hierarchy index, eccentricity (Table 10) and receptive field size (Table 11) were significant. Right V3 showed the highest correlation between eccentricity and hierarchy index at 0.368 whereas left V4 showed the highest correlation between hierarchy index and receptive field size at 0.236. The hierarchy index is calculated for each vertex for each run for each subject and then averaged across runs. The resulting hierarchy index for each vertex is correlated across vertices with the pRF measurement (see Methods). For eccentricity, only the third and fourth runs of the retinotopic data were included in this analysis, since those runs depicted expanding and contracting rings meant to elicit high vs low eccentricity contrasts (see Methods). Lower but statistically significant correlations were observed when all runs were included as well (see Supplementary Fig. S4 A-B). For receptive field size, all retinotopic runs were included. Distributions of these correlations across subjects for the right hemisphere are illustrated in Figure 15 B-C (boxes = median±IQR, gray dots = individual subjects; for the left hemisphere, see Supplementary Fig. S4 C-D).

**Figure 15:**
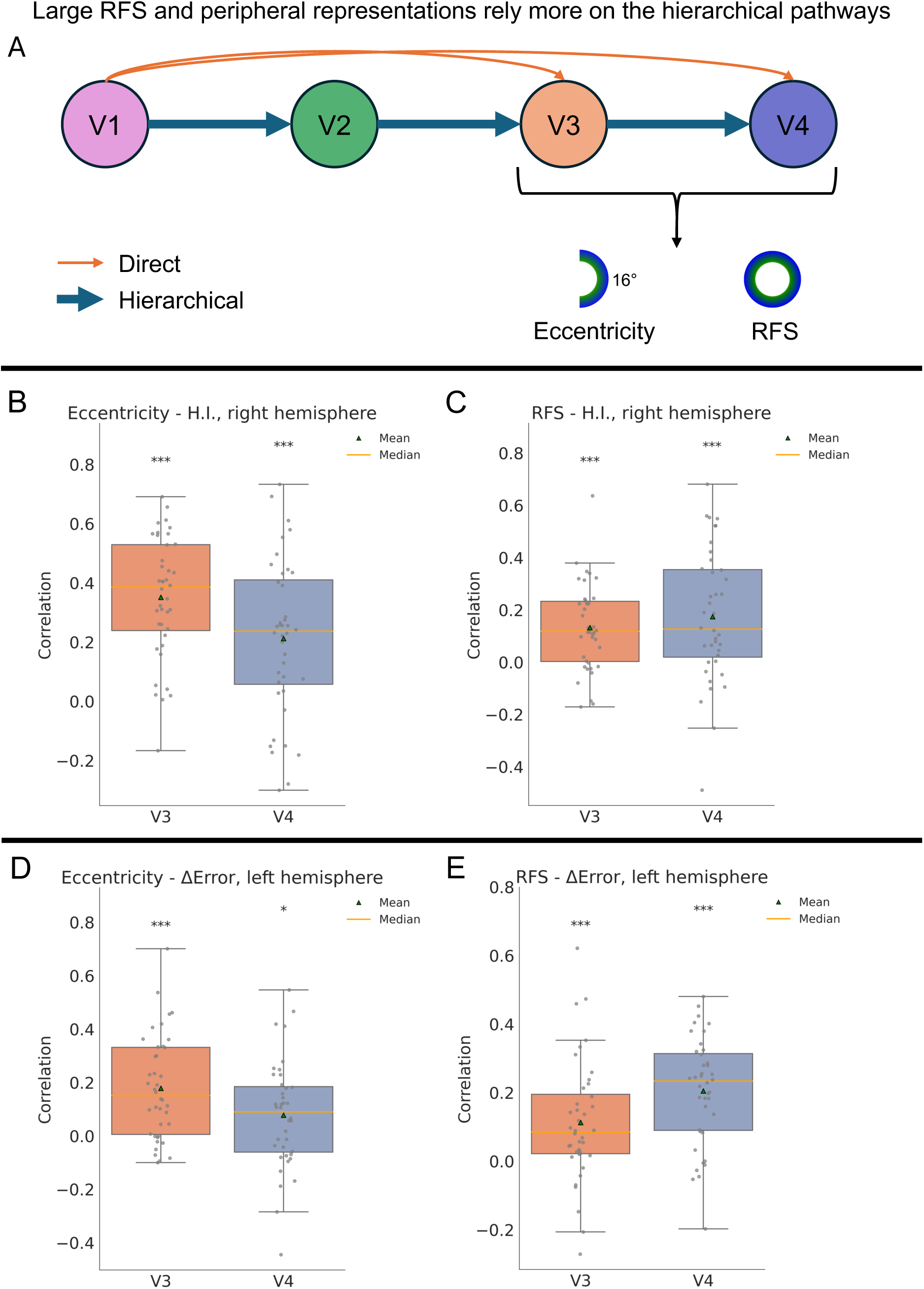
Large receptive fields and peripheral representations rely more on the hierarchical pathways. A: Illustration of the hypothesis that higher eccentricity and receptive field size representations in V3-V4 rely more on the hierarchical pathways. B: Correlations between eccentricity values and hierarchy index for the right hemisphere. All correlations are significant (p<0.0001) (Table 10). The hierarchy index calculated is based on the third and fourth retinotopic runs since these are the runs that feature high vs low eccentricity contrast. C: Correlations between receptive field size and hierarchy index. All runs are included in the calculation of the hierarchy index. All correlations are Bonferroni-corrected for multiple comparisons (Table 11). D: Correlation between eccentricity and the difference of prediction accuracy between the ENN models for the left hemisphere. Larger difference means hierarchical model is better. All correlations are significant (p<0.05) except for right V4 (Table 12). E: Correlation between receptive field size and the difference of prediction accuracy between the ENN models for the left hemisphere. All correlations are significant (p<0.01) (Table 13). For contralateral hemisphere results, see Supplementary Fig. S4. For replication cohort, see Supplementary Fig. S11. Asterisks denote significance: *=p<0.05, **=p<0.01, ***=p<0.001.

**Table 10.** Summary of statistics for correlation between hierarchy index and eccentricity.

| Eccentricity | Correlation | t(39) | Cohen's d | P - values |
| --- | --- | --- | --- | --- |
| Left V3 | 0.306 | 7.349 | 1.162 | p < 0.0001 |
| Left V4 | 0.324 | 9.496 | 1.501 | p < 0.0001 |
| Right V3 | 0.368 | 10.205 | 1.614 | p < 0.0001 |
| Right V4 | 0.230 | 4.985 | 0.788 | p < 0.0001 |

Peripheral representations (higher eccentricity) are significantly correlated with higher values of the hierarchy index. One-sample, one-sided t-tests (mean > 0) are applied. P-values are Bonferroni corrected for multiple comparisons.

**Table 11.** Summary of statistics for correlation between hierarchy index and receptive field size.

| RFS | Correlation | t(39) | Cohen's d | P - values |
| --- | --- | --- | --- | --- |
| Left V3 | 0.082 | 3.458 | 0.547 | p = 0.0027 |
| Left V4 | 0.236 | 6.574 | 1.039 | p < 0.0001 |
| Right V3 | 0.138 | 4.910 | 0.776 | p < 0.0001 |
| Right V4 | 0.188 | 4.298 | 0.679 | p = 0.0002 |

Increased receptive field sizes are significantly correlated with higher values of the hierarchy index. One-sample, one-sided t-tests (mean > 0) are applied. P-values are Bonferroni corrected for multiple comparisons.

**Table 12.**
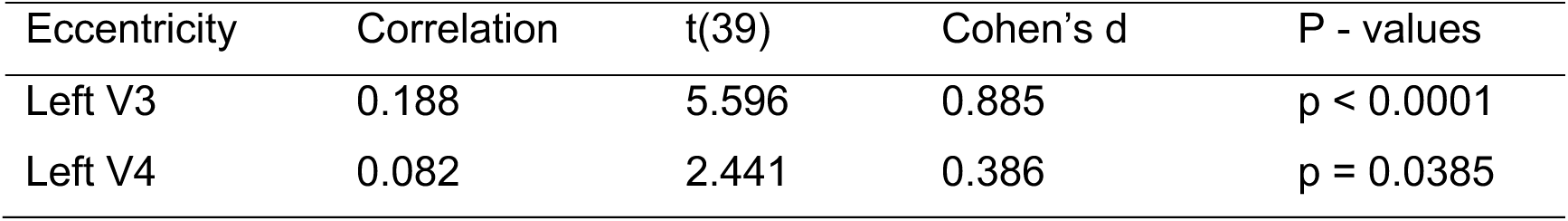
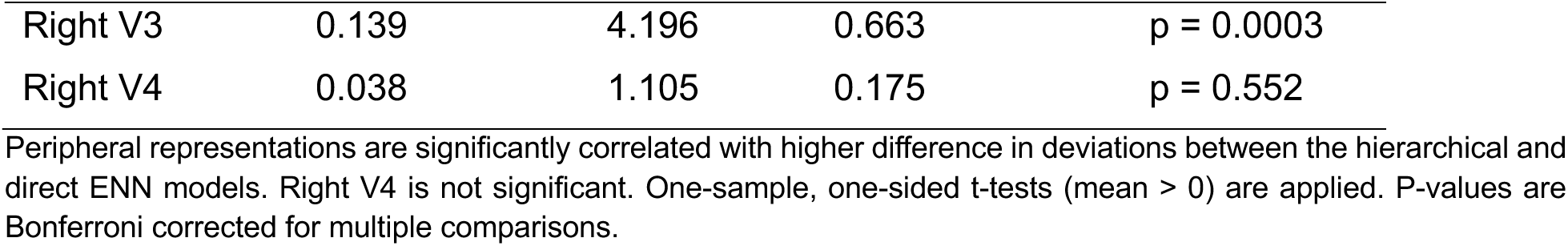
Summary of statistics for correlation between difference of deviations of the hierarchical and direct ENN models and eccentricity.

| Eccentricity | Correlation | t(39) | Cohen's d | P - values |
| --- | --- | --- | --- | --- |
| Left V3 | 0.188 | 5.596 | 0.885 | $p < 0.0001$ |
| Left V4 | 0.082 | 2.441 | 0.386 | $p = 0.0385$ |
| Right V3 | 0.139 | 4.196 | 0.663 | p = 0.0003 |
| Right V4 | 0.038 | 1.105 | 0.175 | p = 0.552 |

Peripheral representations are significantly correlated with higher difference in deviations between the hierarchical and direct ENN models. Right V4 is not significant. One-sample, one-sided t-tests (mean > 0) are applied. P-values are Bonferroni corrected for multiple comparisons.

**Table 13.**
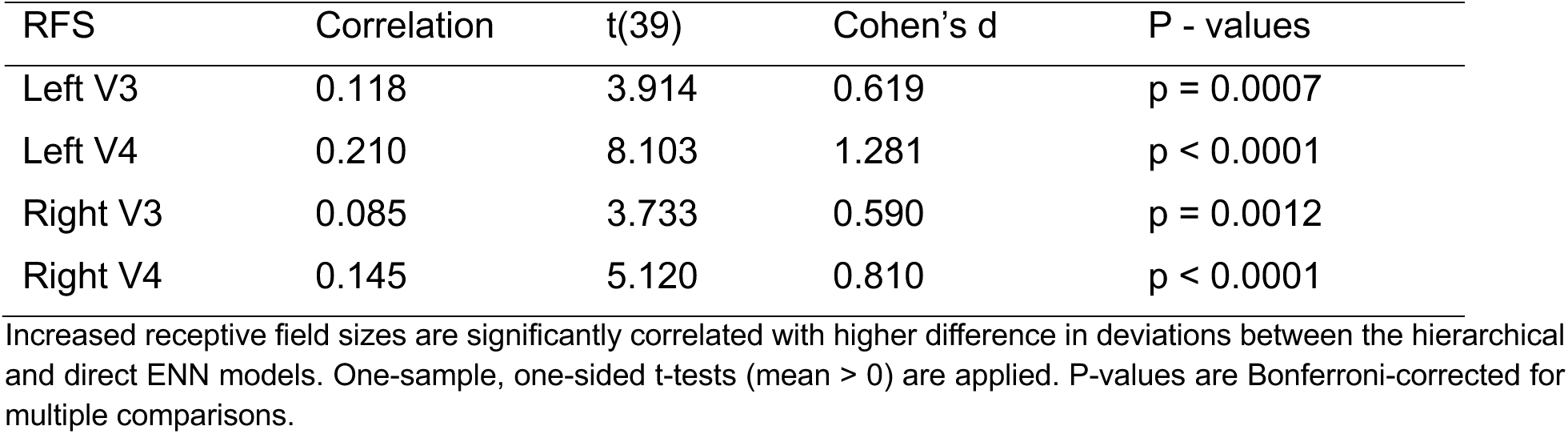
Summary of statistics for correlation between difference of deviations of the hierarchical and direct ENN models and receptive field size.

| RFS | Correlation | t(39) | Cohen's d | P - values |
| --- | --- | --- | --- | --- |
| Left V3 | 0.118 | 3.914 | 0.619 | p = 0.0007 |
| Left V4 | 0.210 | 8.103 | 1.281 | p < 0.0001 |
| Right V3 | 0.085 | 3.733 | 0.590 | p = 0.0012 |
| Right V4 | 0.145 | 5.120 | 0.810 | p < 0.0001 |

Increased receptive field sizes are significantly correlated with higher difference in deviations between the hierarchical and direct ENN models. One-sample, one-sided t-tests (mean > 0) are applied. P-values are Bonferroni-corrected for multiple comparisons.

Second, while these vertex-wise correlations establish an association, they do not test whether hierarchical flows generate larger receptive field sizes and peripheral representations relative to non-hierarchical (direct) flows. We therefore performed a complementary *in silico* experiment. We provided both the hierarchical and the direct activations of the last ENN step to the pRF model and generated retinotopic measurements. For each vertex we calculated the difference for eccentricity and receptive field size accuracies between the two models. Larger difference meant that the hierarchical model was closer to the actual values compared to the direct. We then correlated those differences with the actual eccentricity and receptive field size values for each parcel (see Methods).

Consistent with our hypothesis and the previous finding, we found a positive correlation between the hierarchical vs. direct pathways difference and the actual pRF measurements, further corroborating the finding that large receptive field sizes and peripheral representations rely more on the hierarchical pathways. The highest correlations were observed for left V3 (0.188) for eccentricity and for left V4 (0.210) for receptive field size. All correlations across subjects were significant (Bonferroni corrected for multiple comparisons) except for right V4 for eccentricity (Table 12 and Table 13). Distributions of these correlations across subjects for the left hemisphere are illustrated in Figure 15 D-E (For the left hemisphere, see Supplementary Fig. S4 C-D. For replication cohort, see Supplementary Fig. S11). Angle did not show a significant correlation for any of the regions for either of the analyses, demonstrating that angle representations are encoded irrespective of the pathways.

## Discussion

Hierarchy is often treated as a central organizing principle of the visual system, yet its implications remain unsettled because direct cortico-cortical connections complicate a purely serial V1↔V2↔V3↔V4 account. This ambiguity makes it difficult to determine not only whether the early visual system is hierarchical in practice, but also what distinct functional roles hierarchical and direct pathways play in generating visual functionality. To address this, we adopted two independent operational definitions of hierarchy: a network-based measure derived from a confound-controlled functional connectome and a representational measure derived from stimulus-evoked retinotopic responses. These measures converged on the canonical V1↔V2↔V3↔V4 ordering, providing an empirical scaffold for dissociating pathway contributions. Building on that scaffold, a V1-initiated ENN showed that both hierarchical and direct routes contribute to retinotopic function, but with complementary roles elaborated below.

We began with the network-based definition and our choice of communicability as a flow-sensitive measure of hierarchical distance. Unlike shortest-path metrics, communicability incorporates all possible paths through the network, making it well suited for a system in which direct and multi-step routes coexist. Applied to our confound-controlled GC-PCR connectome, it estimated hierarchical distance in terms of how easily activity could propagate from V1 while reducing inflation of spurious direct connections that would otherwise bias the system toward appearing less hierarchical. This type of inflation is a known issue for the field-standard RSFC measure of pairwise Pearson correlation (Peterson et al., 2025; Sanchez-Romero & Cole, 2021). Another potential issue for our estimates is that RSFC can be inflated at short cortical distances. However, it has been shown that measurements based on partial correlation (as in our GC-PCR FC approach) help control these effects of cortical distance (Dawson et al., 2016).

Representational distance captured hierarchical distance in functional terms, quantifying how much task-evoked retinotopic representations diverged from V1. Though representational similarity analysis has been used to delineate the functional hierarchy in terms of dissimilarity (Gifford et al., 2025), by basing this measure on retinotopic activation dynamics rather than static similarity alone, it better reflects the temporally evolving nature of visual processing in early visual cortex. The convergence of these two independent measures on the canonical V1↔V2↔V3↔V4 ordering strengthens the conclusion that early visual cortex exhibits a prominent hierarchical organization and provides a scaffold for dissociating the functional roles of hierarchical and direct pathways.

Furthermore, the V1-initiated ENN generated retinotopic organization in V2–V4, supporting the conclusion that empirically estimated connectivity, combined with V1 activity, is sufficient to generate retinotopic function throughout early visual cortex. Because the functional connectome provides the scaffold over which activity flows propagate, GC-PCR offered an empirical basis for these generative inferences, while group-average SC increased confidence that the modeled direct V1–V4 connections reflect biologically plausible routes.

This generative framework goes beyond descriptive approaches that link RSFC to retinotopy, such as connectopic mapping (Haak et al., 2018) and connective field modeling (Haak et al., 2013), by explicitly generating task-evoked responses in target regions. Relative to prior V1-initiated ENN work showing that activity flows can generate category selectivity from time-averaged responses (Cocuzza et al., 2024), the present approach used the full fMRI time series rather than time-averaged activations, enabling the generation of retinotopic measurements and the *in silico* dissociation of hierarchical and direct pathway contributions.

We found that both hierarchical and direct pathways contribute to visual function, supporting a view of early visual cortex in which hierarchical processing coexists with parallel routes of information propagation. This is broadly consistent with heterarchical accounts of visual organization (Hawkins et al., 2025), but our results go beyond that general claim by dissociating the mechanistic roles of distinct pathway classes. Unlike most DNN-brain comparisons, where architectural components do not map cleanly onto anatomically defined routes, our empirically constrained ENN enabled targeted *in silico* lesions of specific biological pathways and therefore allowed pathway-specific interpretation. More broadly, although the ENN resembles a DNN in implementing staged transformations of input activity, it differs in several critical respects: its weights and inputs are empirically observed rather than learned to optimize an external objective, it includes feedback interactions, and its topology is not forced into a strictly feedforward hierarchy.

This dissociation revealed complementary representational roles. Hierarchical pathways compressed visual information, whereas direct pathways preserved more high-dimensional representations from V1. This pattern is consistent with broader evidence that dimensionality decreases along the visual hierarchy (Sorscher et al., 2022) and with theoretical work suggesting that deeper processing stages promote integration while direct routes preserve input detail (Furusho & Ikeda, 2020; Singer, 2014). Functionally, lower-dimensional representations may support robustness and generalization, whereas higher-dimensional representations may preserve encoding capacity (Pavuluri & Kohn, 2024). Applied to our findings, this suggests that hierarchical pathways support gradual signal integration into more stable, generalizable representations, whereas direct pathways preserve richer detail and faster access to visual information. In this sense, the two pathway classes may support a tradeoff between robustness and generalization on the one hand and encoding capacity and fine-grained visual detail on the other. Dimensionality was quantified using the participation ratio computed from cross-validated neural representations, reducing noise inflation. Although parcel size can influence dimensionality estimates, it is also a stable characteristic of the system, rather than a nuisance (Eliasmith, 2013). In addition, our critical dimensionality comparisons were made between models rather than between parcels, thereby controlling this factor across pathway-lesion conditions.

Lastly, we demonstrated that larger receptive field sizes and peripheral representations rely more on the hierarchical pathways, providing important additional detail regarding the role of hierarchy in generating visual functionality. To our knowledge, this is the first evidence that dissociates routing by eccentricity in the human brain. This eccentricity-dependent dissociation is in accordance with macaque studies (Nakamura et al., 1993; Yukie & Iwai, 1985) that demonstrate direct V1 to V4 foveal connections. In humans, the existence of direct connections favoring central visual representations is also evident for functional and anatomical connections between V1 and frontal regions (Sims et al., 2021).

Prior literature demonstrated that direct links are utilized by the visual system to rapidly alert higher order regions in the dorsal stream about motion, potentially offering an evolutionary advantage (Laycock et al., 2007; Sincich et al., 2004). Direct pathways feature decreased latency due to less intermediate layers of information processing, rendering them appropriate for fast motion detection. This does not contradict the need for hierarchical construction of large receptive field sizes in early visual cortex. Rather, it complements our finding by distinguishing alerting speed from multi-step construction of representations.

Several future directions follow from the present work. First, spatial confounding remains an important consideration for activity-flow analyses in visual cortex, where source and target vertices can be anatomically close (Watson & Andrews, 2023). However, Aquino and colleagues demonstrated that outside of a 5 mm radius, spatial autocorrelation becomes negligible (Aquino et al., 2012). Consistent with this, prior activity-flow studies have often used 10 mm exclusion zones to reduce such effects, but we did not apply that dilation here because short-range connections are especially prominent in visual cortex, with 80-90% of connections residing within 1–2 mm of each potential target (Vezoli et al., 2021). Importantly, a control analysis with a 5 mm exclusion area produced only a negligible drop in explained variance (Supplementary Fig. S5), suggesting that our main findings are not driven by local spatial leakage.

A second important direction is to extend this framework beyond corticocortical interactions alone. Subcortical structures such as the lateral geniculate nucleus and pulvinar may play important roles in shaping both hierarchical and direct routes, complicating any purely cortical account of visual hierarchy (Bridge et al., 2016; Cortes et al., 2023). Future work should also test whether the pathway dissociations observed here extend to higher-order visual cortex and behavior—for example, whether hierarchical routes preferentially support peripheral integration and scene-level processing, whereas direct routes support foveally detailed object information and faster responses. More mechanistically, feedback lesions could clarify how recurrent interactions contribute to hierarchical compression, and the hierarchy index introduced here may prove useful as a biomarker of visual pathway integrity.

To conclude, utilizing high-resolution 7T fMRI data collected during rest and retinotopic mapping along with diffusion MRI data, we demonstrated that the early visual system implements a dual-route policy: hierarchical paths provide compressed, generalizable representations whereas direct paths deliver rapid, detail-rich representations. The visual system uses activity flows to flexibly blend these representational features when generating visual functionality, helping to reconcile the hierarchical versus direct pathways debate in early visual cortex.

## Acknowledgments

This work was supported by the US National Institutes of Health (NIH) grant R56MH138448 and the US National Science Foundation (NSF) under awards 2219323 and 2117429. HCP data were provided by the Human Connectome Project, WU-Minn Consortium (Principal Investigators: David Van Essen and Kamil Ugurbil; 1U54MH091657) funded by the 16 NIH Institutes and Centers that support the NIH Blueprint for Neuroscience Research; and by the McDonnell Center for Systems Neuroscience at Washington University. We thank the Office of Advanced Research Computing at Rutgers, the State University of New Jersey, for providing access to the Amarel cluster and associated research computing resources that have contributed to the results reported here. We thank Dr. Bart Krekelberg and Dr. Brian Keane for detailed feedback on the manuscript and members of the Cole Neurocognition Lab for helpful discussions. The content is solely the responsibility of the authors and does not necessarily represent the official views of any of the funding agencies.

## Data availability

Data analyzed in this study are available from the HCP 7T retinotopy resources described by Benson et al. (2018), subject to HCP data use terms. The data correspond to the HCP-Young Adult 2025 release, accessible via ConnectomeDB powered by BALSA (https://balsa.wustl.edu/project?project=HCP_YA).

## Code availability

Custom code used for the analyses in this study will be made publicly available upon publication.

## Author contributions

A.T. and M.W.C. designed research. A.T. performed research and analyzed data. D.E.O., C.C., L.N.C., R.D.M. and K.L.P. contributed unpublished analytic tools. A.T. wrote the original draft of the manuscript. A.T. and M.W.C. wrote the paper. A.T., D.E.O., C.C., L.N.C., R.D.M., K.L.P. and M.W.C. edited the paper. All authors have approved the final content. M.W.C acquired funding.

