## Supplementary Information for "Hierarchical and non-hierarchical network flows generate complementary representational dynamics in human visual cortex"

### Supplement

#### S1. Additional results – contralateral hemisphere

##### S1.1. Functional connectivity in the left hemisphere

The main results focused on the right hemisphere for conciseness. We report the left hemisphere results here (Supplementary Fig. S1). For the left hemisphere, 100% of subjects featured all the hierarchical connections, 77.5% featured the direct V1-V3 connection, 80% the direct V1-V4 and 80% V2-V4. Percentages were calculated by taking into consideration either of the contralateral sources.

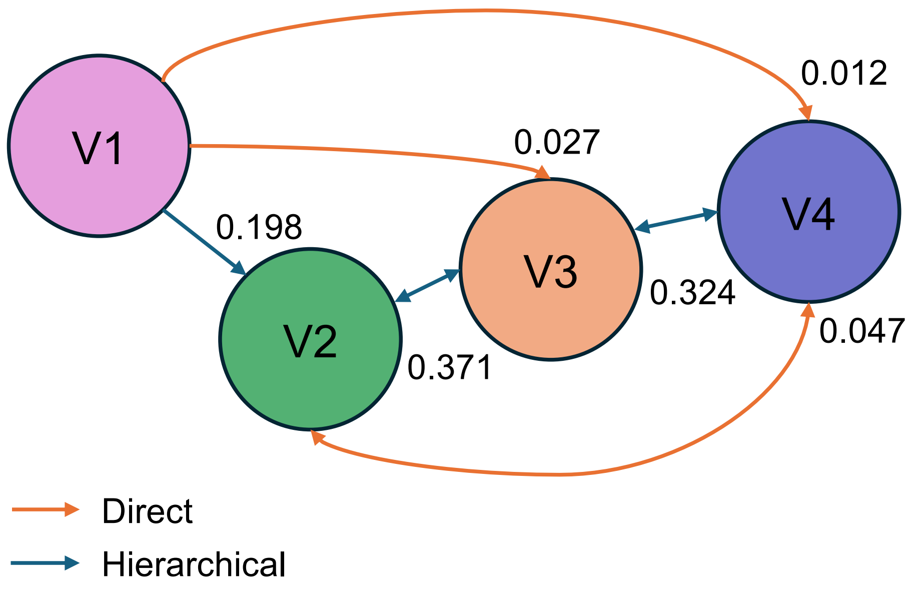


**Supplementary figure S1: Parcel-level connectivity between regions V1-V4 in the left hemisphere (related to Fig. 5C)**. Force-directed graph based on parcel level connectivity for regions left V1-V4 (the results in the main text focused on the right hemisphere). Functional connections are included after masking by group-average structural connectivity (all connections survived). Distances between nodes represent connectivity strength. Shorter distances represent stronger connections. The strongest connection is between V2-V3 and the weakest is between V1-V4. Numbers denote the cross-subject average connectivity strength.

##### S1.2. Network flow distance in the left hemisphere

A similar trend to the right hemisphere (Fig. 6) was observed in the left hemisphere as well for network flow distance. All pairwise differences were statistically significant except for V3-V4 for the GC-PCR based communicability (Supplementary Fig. S2).

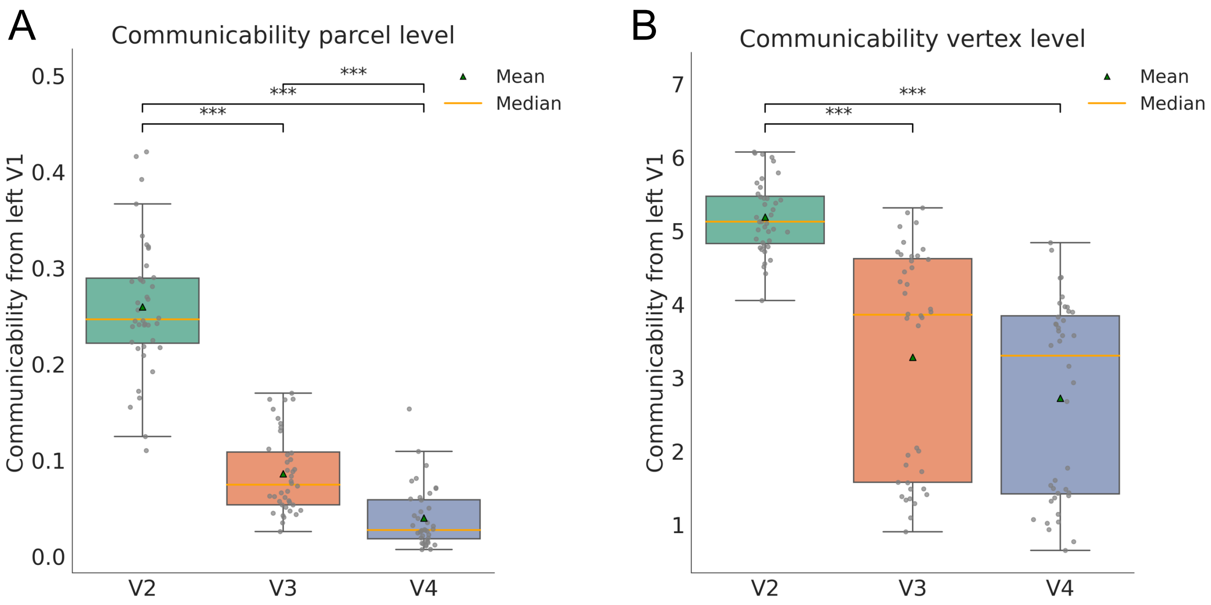


**Supplementary figure S2: Connectivity-based hierarchy for the left hemisphere (related to Fig. 6B-C)**. A: Communicability results based on the parcel level FC graph for the left hemisphere. All pair differences are significant. B: Communicability results based on the vertex level FC graph for the left hemisphere. All pair differences are significant, except for V3-V4. Values are multiplied by $10^4$ for visualization (vertex FC weights ~ $10^{-4}$). Boxes = median±IQR, gray dots = individual subjects. Asterisks denote significance: *=p<0.05, **=p<0.01, ***=p<0.001.

#### S1.3. RSMs for the right hemisphere

Similar to Figure 7, we present the RSMs for the right hemisphere in Supplementary Fig. S3.

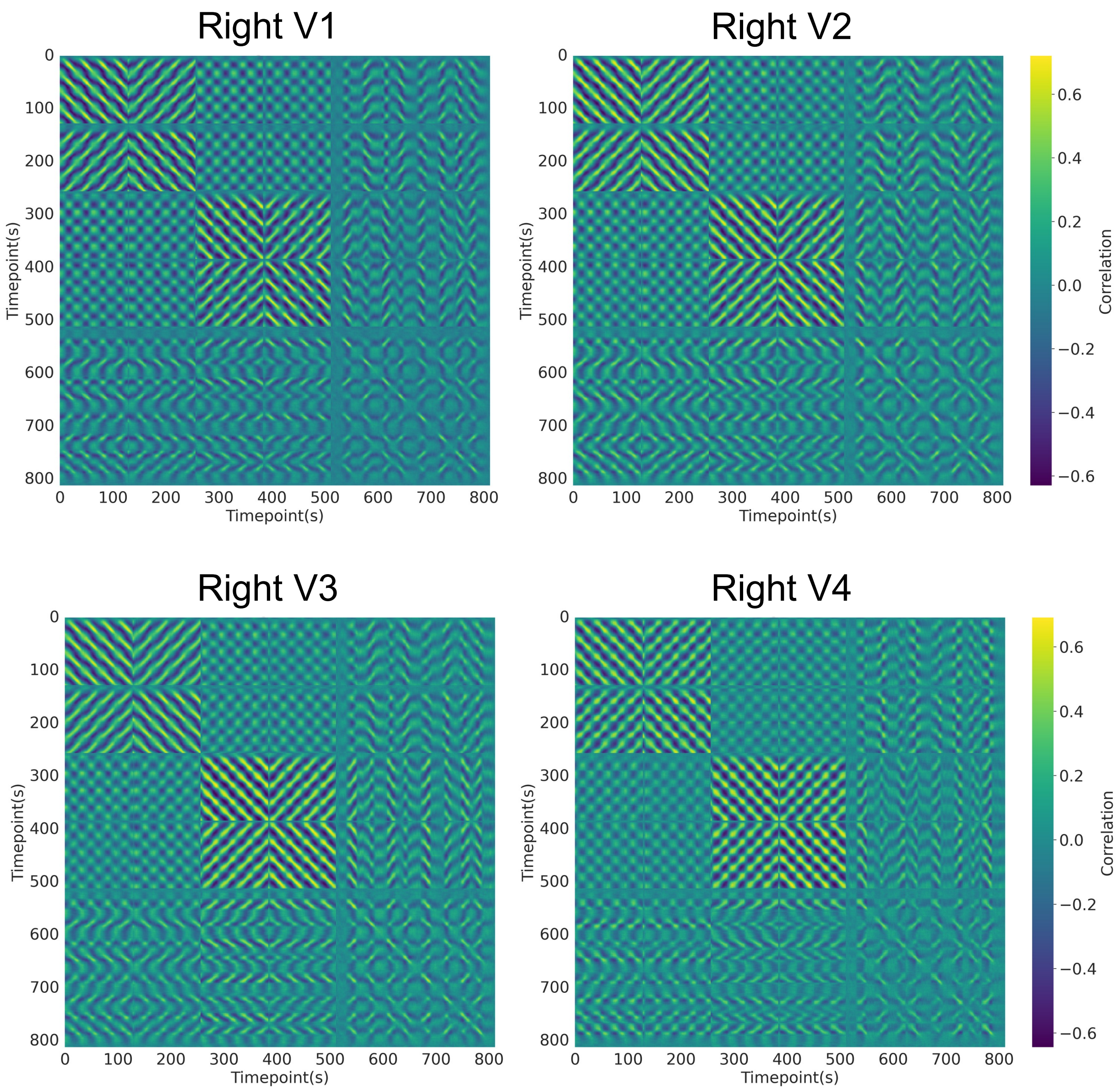


**Supplementary figure S3: Representational similarity matrices for neural time series (related to Fig. 7C)**. Cross-validated RSMs for each region of the early visual system for the right hemisphere.

#### S1.4. Hierarchical pathways and large receptive fields

We present the results when all runs are included in the calculation of the hierarchy index (HI) and in its correlation with eccentricity. Correlation is lower but still significant except for right V4. We also present the correlations between HI and eccentricity and receptive field size for the left hemisphere along with the correlation between the hierarchical-direct deviation and eccentricity and receptive field size for the right hemisphere (Supplementary Fig. S4).

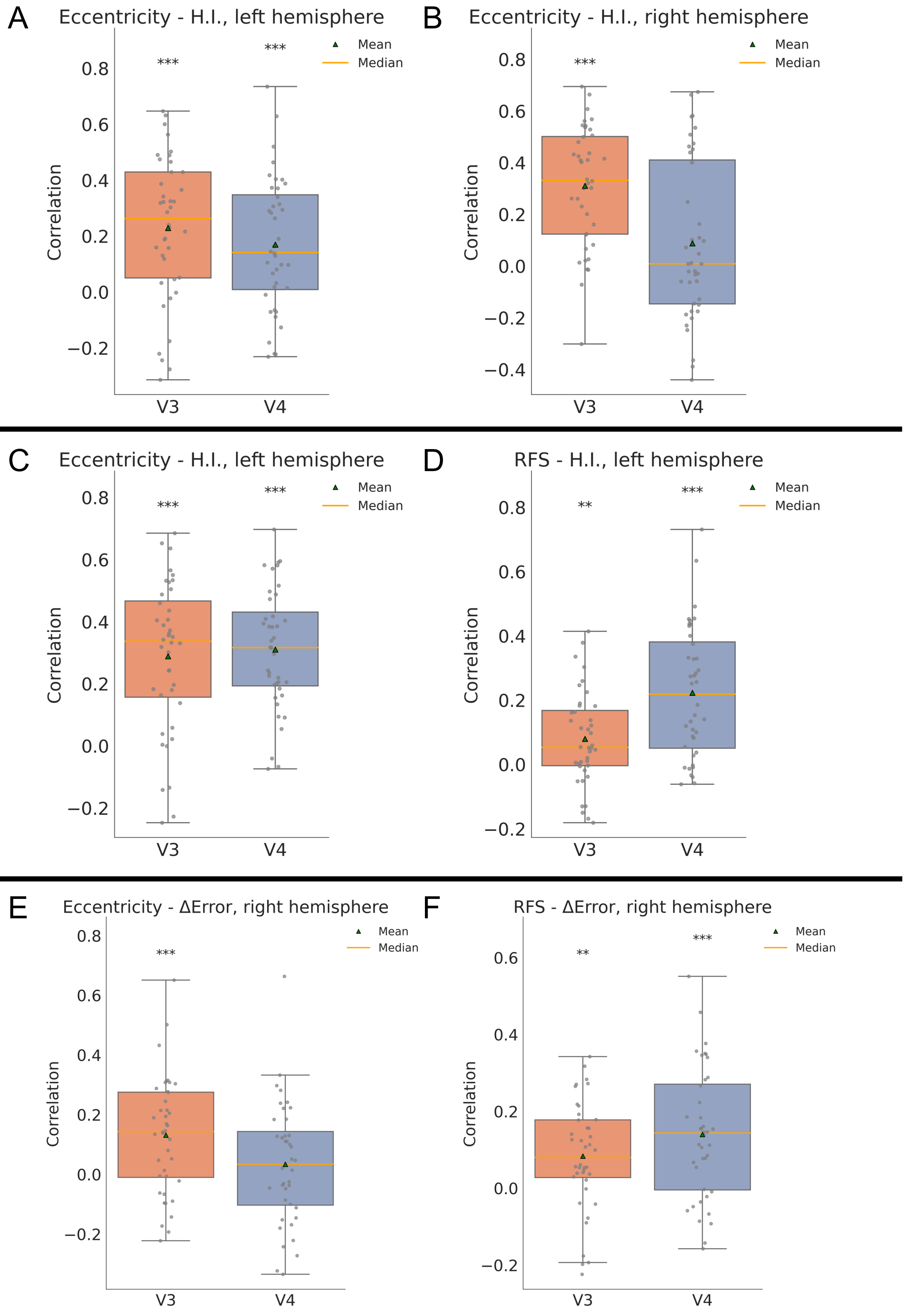
A
Eccentricity - H.I., left hemisphere
B
Eccentricity - H.I., right hemisphere
C
Eccentricity - H.I., left hemisphere
D
RFS - H.I., left hemisphere
E
Eccentricity - ΔError, right hemisphere
F
RFS - ΔError, right hemisphere
Mean
Median
Correlation
V3
V4
***
**
0.8
0.6
0.4
0.2
0.0
−0.2
−0.4

**Supplementary figure S4: The relationship of large receptive fields and peripheral representations with hierarchy for the contralateral hemispheres (related to Fig. 15B-E)**. A-B: Correlation between eccentricity and hierarchy index for left and right hemisphere when all retinotopic runs are included in the estimation of HI. We observe significant results for all regions as in the main analysis, except for right V4. C-D: Correlation between eccentricity and receptive field size and HI for the left hemisphere. All regions show significant correlations. E-F: Correlation between eccentricity and receptive field size and the difference of prediction accuracies between the lesion models for the right hemisphere. All correlations are significant except for right V4 for eccentricity. Boxes = median±IQR, gray dots = individual subjects. Asterisks denote significance: *=p<0.05, **=p<0.01, ***=p<0.001.

### S2. Control Analyses – Main Sample

#### S2.1. Dilation control analysis

During GC-PCR, we applied dilation (an exclusion area around target vertices; see Discussion) on the resting-state fMRI data based on a 5-mm-radius sphere for each target vertex. We used 15 subjects (9 females) from the main sample. For this subset, the original (no-dilation) model converged at 5 steps instead of 6. In supplementary figure S5, V1-initiated ENN model performance across steps is displayed. Explained variance is decreased from the full model (41% vs. 35% variance explained), as expected. However, the minimal decrease in overall variance explained demonstrates that, even if spatial smoothness in the fMRI data inflated activity flow estimation accuracy, that inflation was small.

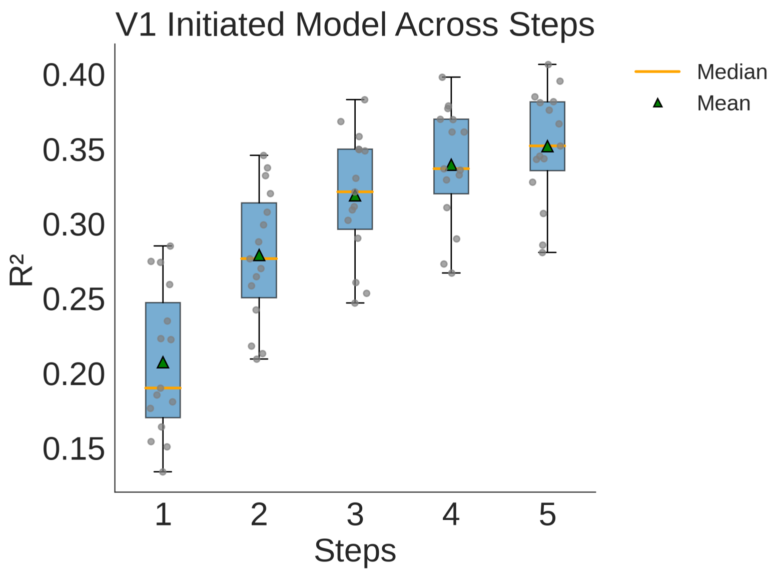


**Supplementary figure S5: V1-ENN model performance for 15 subjects after 5mm dilation (related to Fig. 10A)**. $R^2$ values for each step of the V1-ENN, averaged across runs and parcels, after applying 5 mm dilation. $R^2$ was calculated based on the averaged across vertices time series for each parcel. Averaged across subjects values: 5$^{th}$-step $R^2$ = 35.18%, 4$^{th}$-step $R^2$ = 33.94%, 3$^{rd}$-step $R^2$ = 31.89%, 2$^{nd}$-step $R^2$ = 27.89%, 1$^{st}$-step $R^2$ = 20.74%. Boxes = median±IQR, gray dots = individual subjects.

#### S2.2. Extra lesion analysis

As an additional test, we included one extra lesion for our *in silico* lesion models. For the hierarchical model, we included the direct V2-V4 lesion, whereas for the direct model, we included the hierarchical V1-V2 lesion. This resulted in a hierarchical model with lesions: V1-V3, V1-V4, V2-V4 and direct model with lesions: V1-V2, V2-V3, V3-V4. In supplementary figure S6 we present the contributions of these alternative hierarchical vs. direct models.

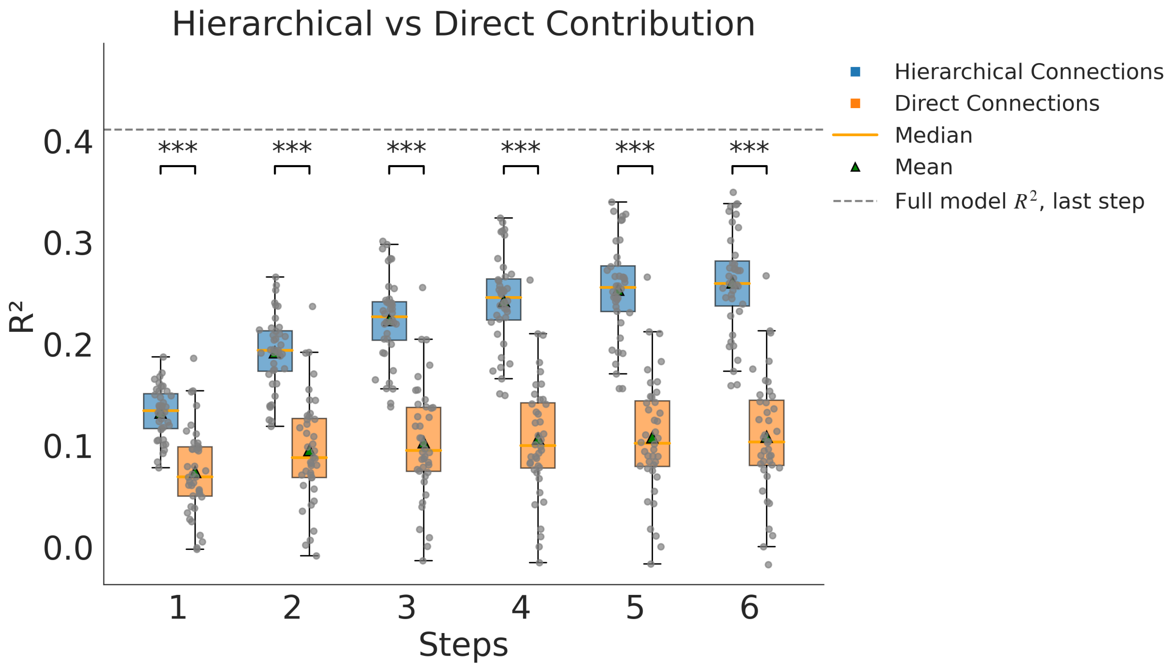


**Supplementary figure S6: Hierarchical vs. direct contributions when V2-V4 and V1-V2 lesions are added (related to Fig. 12B)**. Similar trend as the original lesion models with the hierarchical showing increasing accuracy as the system reaches its steady state. Direct model plateaus after the first step and remains at a low level of explained variance. Both models perform worse compared to their original counterparts as expected. The direct model shows a significant decline, with performance during the original lesions being higher than the direct during the first steps. $R^2$ of the hierarchical model at the last step is 26.07% (original hierarchical was 30.1%) whereas $R^2$ of the direct is 10.89% (original direct was 28.4%). Boxes = median±IQR, gray dots = individual subjects. Asterisks denote significance: *=p<0.05, **=p<0.01, ***=p<0.001.

### S3. Replication Sample (N = 40)

We randomly selected 40 more subjects (24 female) from the 181 healthy young adults HCP S900 release to run a replication analysis. All analyses for the replication sample mirrored the primary analyses described in Methods. Briefly, these include: i) definition of hierarchy based on network flow distance and representational distance (Supplementary Fig. S7), ii) validation of the V1-ENN model (Supplementary Fig. S8) and its ability to generate pRF measurements (Supplementary Fig. S9), iii) hierarchical vs direct contributions in visual functionality (Supplementary Fig. S10), iv) hierarchical vs direct dimensionality (Supplementary Fig. S10), v) pathway preference for large receptive field and peripheral representations (Supplementary Fig. S11). We also include the percentage of connections present between parcels for both hemispheres: for the left hemisphere, 100% of subjects featured all the hierarchical connections, 87.5% featured the direct V1-V3 connection, 70% the direct V1-V4 and 82.5% V2-V4. For the right hemisphere: 100% of subjects featured all the hierarchical connections, 82.5% featured the direct V1-V3 connection, 72.5% the direct V1-V4 and 65% V2-V4. Percentages were calculated by taking into consideration either of the contralateral sources.

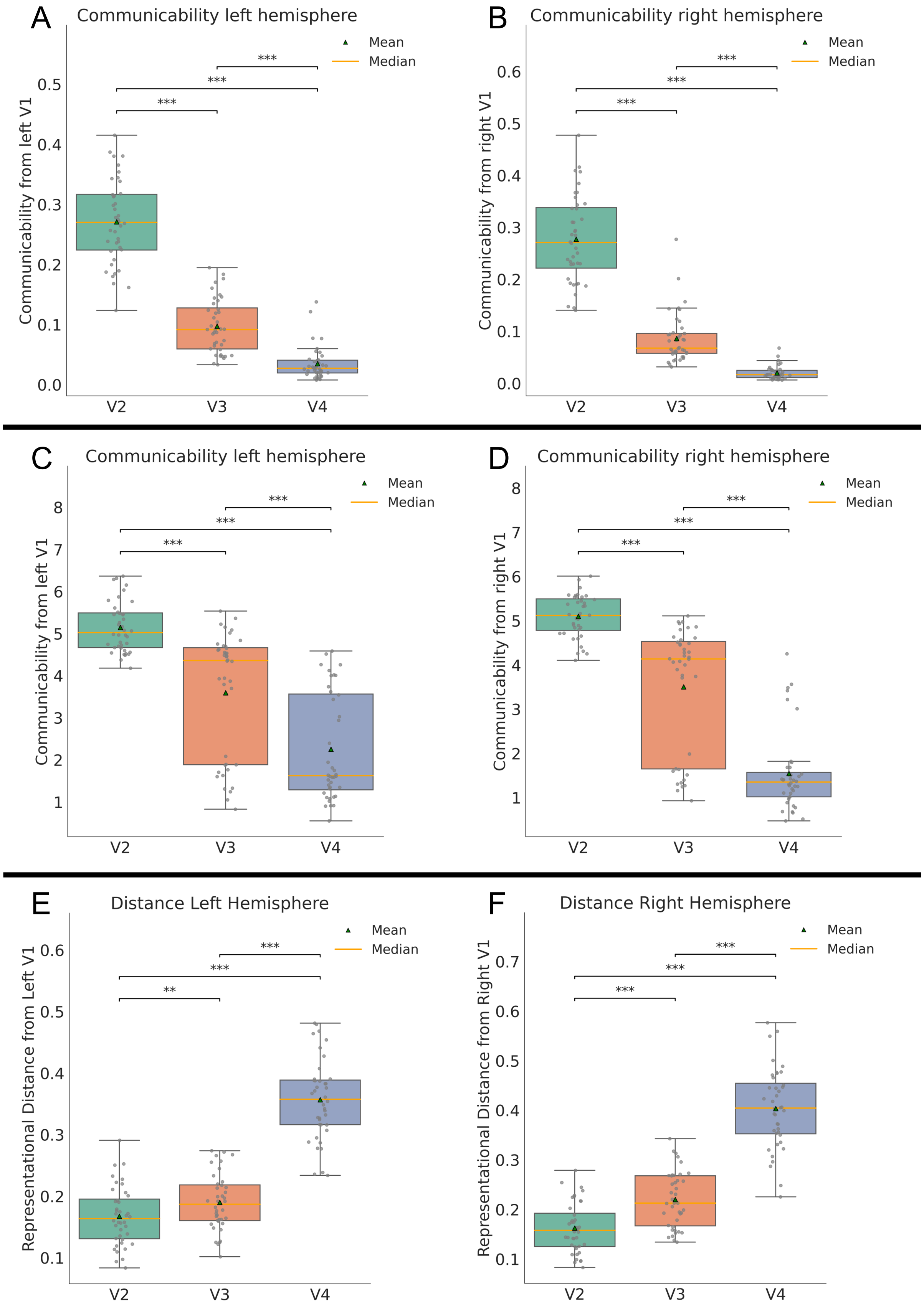

A
Communicability left hemisphere
Communicability from left V1
Mean
Median
***
V2
V3
V4
B
Communicability right hemisphere
Communicability from right V1
C
Communicability left hemisphere
Communicability from left V1
D
Communicability right hemisphere
Communicability from right V1
E
Distance Left Hemisphere
Representational Distance from Left V1
**
F
Distance Right Hemisphere
Representational Distance from Right V1

**Supplementary figure S7: Replication cohort: Hierarchy defined based on network flow distance and representational distance (related to Fig. 6B-C and Fig. 9B)**. A-B: A: Communicability results based on the parcel level FC graph for both hemispheres. All pair differences are significant. C-D: Communicability results based on the vertex level FC graph for both hemispheres. All pair differences are significant. Values are multiplied by $10^4$ for visualization (vertex FC weights ~ $10^{-4}$). E-F: Representational distance for the left and right hemispheres respectively. The pattern is similar across all cases, and the same hierarchy emerges: V1→V2→V3→V4. All region pair comparisons are significant. Boxes = median±IQR, gray dots = individual subjects. Asterisks denote significance: *=p<0.05, **=p<0.01, ***=p<0.001.

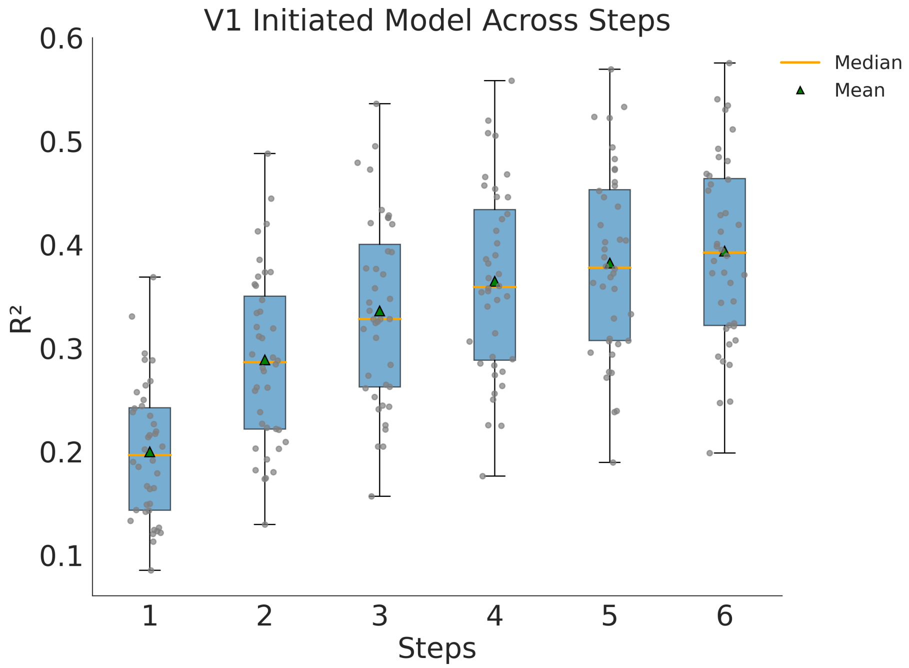


**Supplementary figure S8: Replication cohort: Simulated activity flow using individual-subject FC based ENNs accurately generates retinotopic time series (related to Fig. 10A)**. $R^2$ between predicted and actual time series (across left and right V1, V2, V3, and V4) at each flow step of the model. Calculated after averaging generated and original time series for each parcel. Averaged across retinotopic runs and parcels. Performance plateaus at the 6th step. Averaged values across subjects: step 1 $R^2$ = 19.97%, step 2 $R^2$ = 28.86%, step 3 $R^2$ = 33.60%, step 4 $R^2$ = 36.45%, step 5 $R^2$ = 38.21%, step 6 $R^2$ = 39.36%. Boxes = median±IQR, gray dots = individual subjects.

Supplementary table S1. Replication cohort: Activity flow model retinotopic time series prediction accuracy (parcel level).

| Region | Pearson's r | $R^2$ (%) | P-value |
|---|---|---|---|
| Left V2 | 0.863 | 57.23 | p < 0.0001 |
| Left V3 | 0.808 | 46.99 | p < 0.0001 |
| Left V4 | 0.669 | 24.79 | p < 0.0001 |
| Right V2 | 0.832 | 51.92 | p < 0.0001 |
| Right V3 | 0.764 | 38.60 | p < 0.0001 |
| Right V4 | 0.605 | 16.62 | p < 0.0001 |

Pearson r and $R^2$ were calculated across the average time series for each parcel and for each subject. Here we present the average across subjects. P-values were obtained after running the Wilcoxon test on the $R^2$ distribution across subjects. Bonferroni corrected for multiple comparisons.

Supplementary table S2. Replication cohort: Activity flow model retinotopic time series prediction accuracy (vertex level).

| Region | Pearson's r | $R^2$ (%) | P-value |
|---|---|---|---|
| Left V2 | 0.524 | 20.04 | p < 0.001 |
| Left V3 | 0.518 | 19.23 | p < 0.001 |
| Left V4 | 0.457 | 13.61 | p < 0.001 |
| Right V2 | 0.482 | 16.97 | p < 0.001 |
| Right V3 | 0.493 | 16.39 | p < 0.001 |
| Right V4 | 0.371 | 8.93 | p < 0.001 |

Pearson r and $R^2$ were calculated for each vertex for each subject. Here the average r and $R^2$ across vertices for each region and across subjects is reported. P-values based on $R^2$ are obtained from a MaxT nonparametric permutation test.

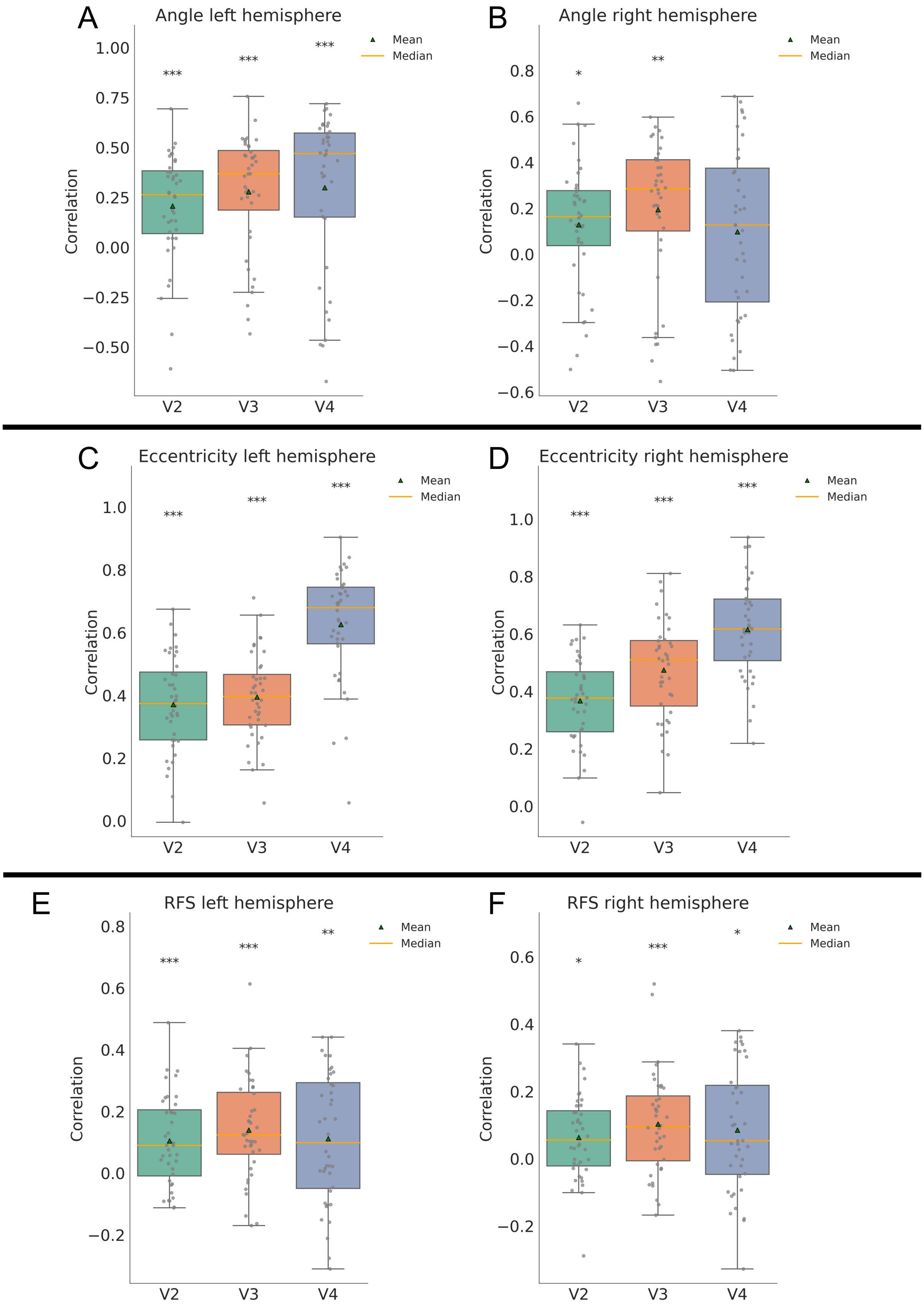

A
Angle left hemisphere
B
Angle right hemisphere
C
Eccentricity left hemisphere
D
Eccentricity right hemisphere
E
RFS left hemisphere
F
RFS right hemisphere
Mean
Median
Correlation
V2
V3
V4

**Supplementary figure S9: Replication cohort: pRF measurements can be accurately generated by simulated flows over FC (related to Fig. 11)**. A-B: Circular correlation between actual and generated angle across vertices for each parcel of the early visual system for left and right hemisphere respectively. Correlations are significant for all parcels ($p < 0.0001$) except for right V4. C-D: Correlations between actual and generated eccentricity for each parcel of the early visual system for left and right hemisphere respectively. Correlations are significant for all parcels ($p < 0.0001$). E-F: Correlations between actual and generated receptive field size for each parcel of the early visual system for left and right hemisphere respectively. Correlations are significant for all parcels ($p < 0.05$). Boxes = median±IQR, gray dots = individual subjects. Asterisks denote significance: *=p<0.05, **=p<0.01, ***=p<0.001.

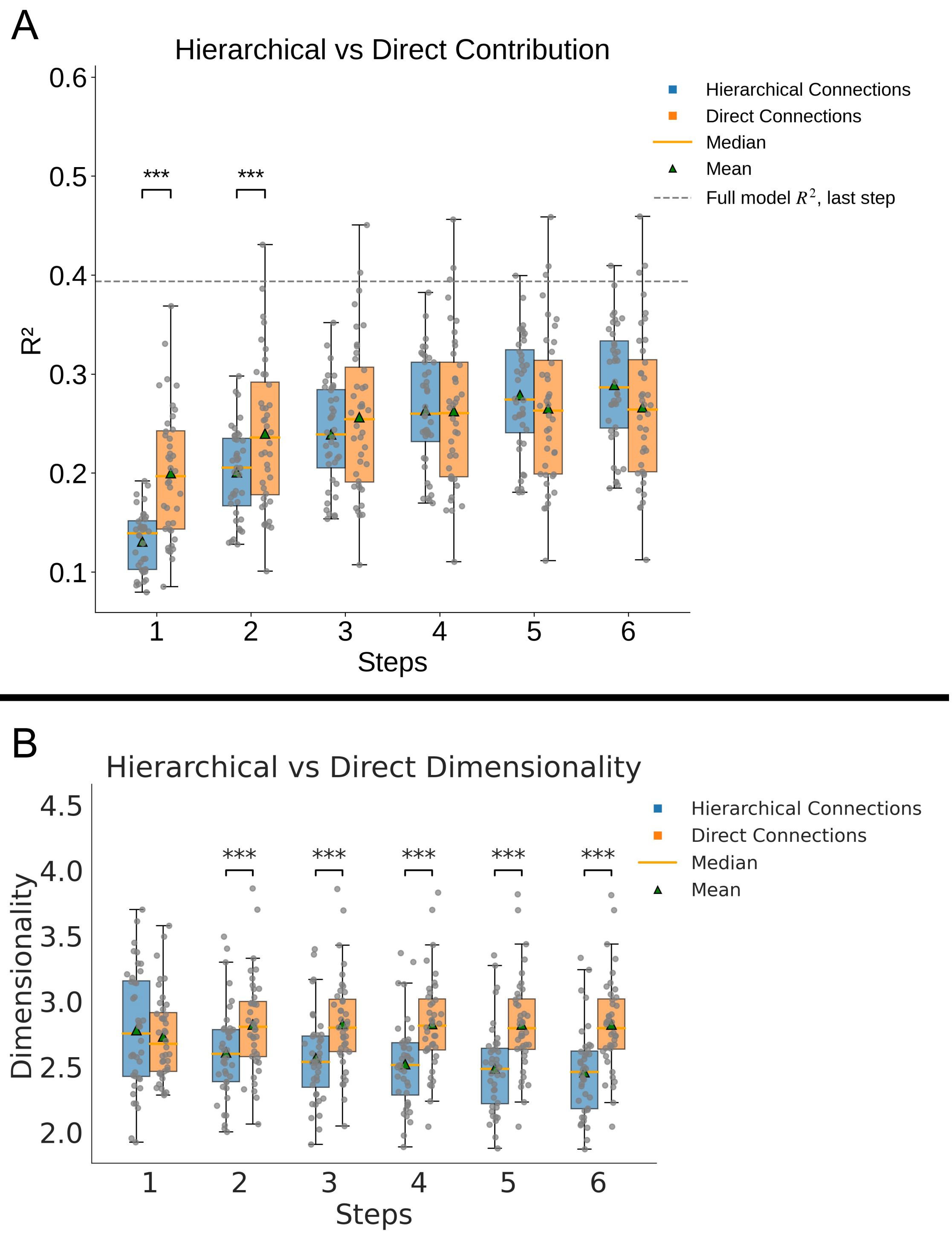


**Supplementary figure S10: Replication cohort: *In silico* ENN lesioning experiment: Both hierarchical and direct connections contribute to visual functionality generation. Hierarchical pathways compress visual representations (related to Fig. 12B and Fig. 14B)**. A: Comparison between the two different lesion models at the vertex level. Generated time series were averaged for each parcel and then compared to the original ones resulting in $R^2$ values. These values were averaged across runs and parcels. Initially, and similar to the first group, the two models are significantly different with the hierarchical pathways demonstrating larger increase across steps, outperforming the direct one at the last two steps (difference not significant). B: Dimensionality across steps for the hierarchical and direct pathways. Excluding the first step, all differences were statistically significant and showed increased dimensionality for the direct pathways. Boxes = median±IQR, gray dots = individual subjects. Asterisks denote significance: *=p<0.05, **=p<0.01, ***=p<0.001.

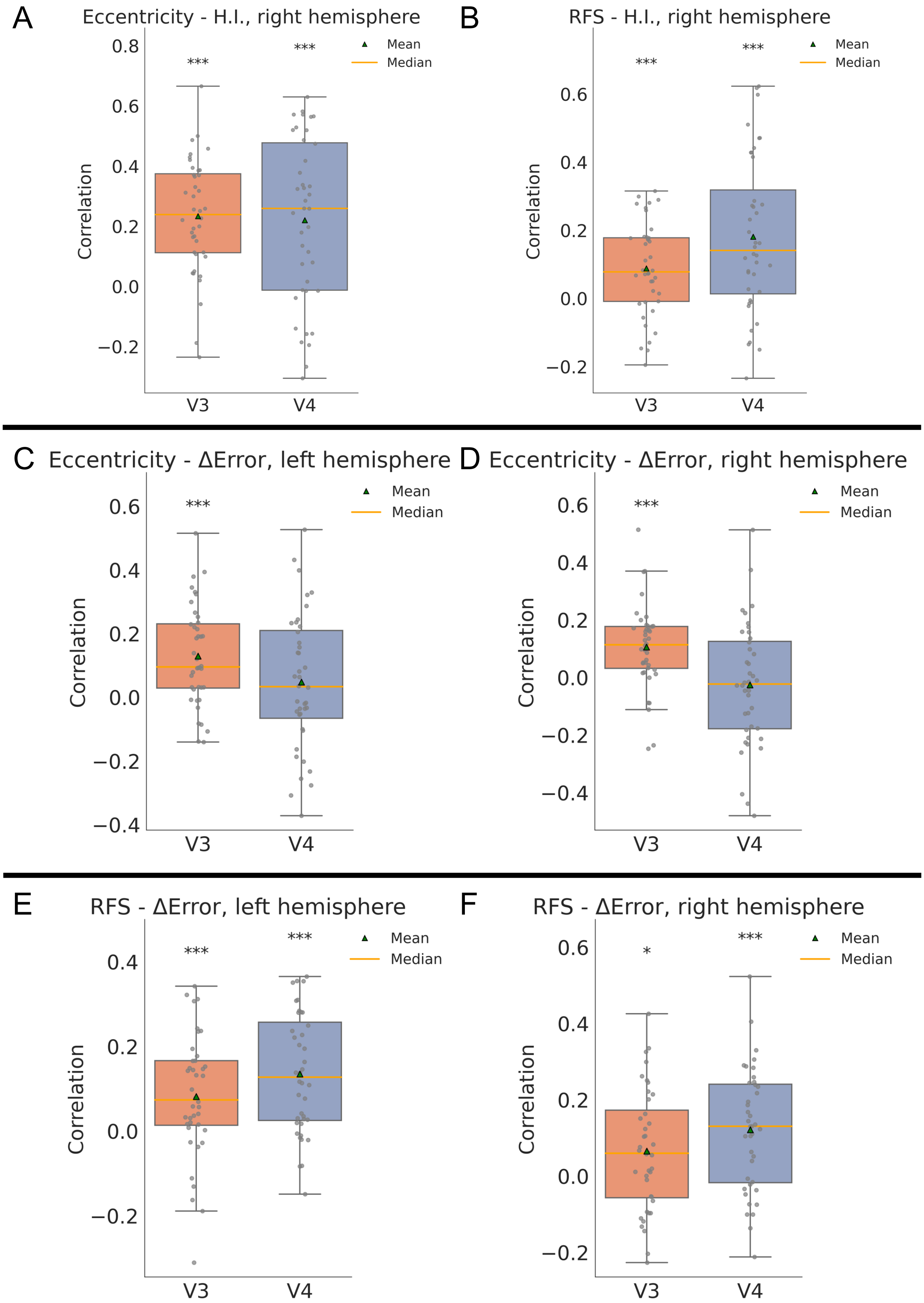
A
Eccentricity - H.I., right hemisphere
Correlation
0.8
0.6
0.4
0.2
0.0
−0.2
V3
V4
***
***
Mean
Median
B
RFS - H.I., right hemisphere
Correlation
0.6
0.4
0.2
0.0
−0.2
V3
V4
***
***
Mean
Median
C
Eccentricity - ΔError, left hemisphere
Correlation
0.6
0.4
0.2
0.0
−0.2
−0.4
V3
V4
***
Mean
Median
D
Eccentricity - ΔError, right hemisphere
Correlation
0.6
0.4
0.2
0.0
−0.2
−0.4
V3
V4
***
Mean
Median
E
RFS - ΔError, left hemisphere
Correlation
0.4
0.2
0.0
−0.2
V3
V4
***
***
Mean
Median
F
RFS - ΔError, right hemisphere
Correlation
0.6
0.4
0.2
0.0
−0.2
V3
V4
*
***
Mean
Median

**Supplementary figure S11: Replication cohort: Large receptive fields and peripheral representations rely more on the hierarchical pathways (related to Fig. 15B-E).** A: Correlations between eccentricity values and hierarchy index for the right hemisphere. All correlations are significant for both left and right hemispheres ($p<0.001$). The hierarchy index calculated is based on the third and fourth retinotopic runs since these are the runs that feature high vs low eccentricity contrast. When all runs were included, correlations dropped but were still significant for all regions except left V4 ($p<0.05$). B: Correlations between receptive field size and hierarchy index for the right hemisphere. All runs are included in the calculation of the hierarchy index. Correlations were significant for both hemispheres ($p<0.01$). C-D: Correlation between eccentricity and the difference in prediction accuracy between the models for left and right hemispheres respectively. A larger difference means that the hierarchical model performed better. Correlations are significant for V3 ($p<0.001$) but not for V4. E-F: Correlation between receptive field size and the difference of prediction accuracy between the models for the left and right hemispheres. All correlations are significant ($p<0.05$). Asterisks denote significance: *=$p<0.05$, **=$p<0.01$, ***=$p<0.001$.
